# Metabolic reshaping drives novel melanin production from tryptophan in a cystic fibrosis *Pseudomonas aeruginosa* clinical isolate

**DOI:** 10.64898/2025.12.15.694361

**Authors:** Mateo N. Diaz Appella, Adriana A. Kolender, Mulugeta Nega, Stefania A. Robaldi, Pablo M. Cassanelli, Ningna Li, Elisa Liberini, Pablo Hoijemberg, Leonardo Pellizza, Martin Aran, Friedrich Götz, Nancy I. López, Paula M. Tribelli

**Affiliations:** Instituto de Química Biológica (IQUIBICEN)-Universidad de Buenos Aires (UBA)-Consejo Nacional de Investigaciones Científicas y Técnicas (CONICET), Buenos Aires, Argentina; Departamento de Química Orgánica, Facultad de Ciencias Exactas y Naturales (FCEyN), UBA, Buenos Aires, Argentina; Centro de Investigaciones en Hidratos de Carbono (CIHIDECAR),UBA-CONICET, Buenos Aires, Argentina; Department of Microbial Genetics, Interfaculty Institute of Microbiology and Infection Medicine Tübingen (IMIT), University of Tübingen, Tübingen, Germany; Hospital General de Niños ―Pedro de Elizalde‖, Buenos Aires, Argentina; Centro de Investigaciones en Bionanociencias (CIBION), CONICET, Godoy Cruz 2390, C1425FQD Ciudad Autónoma de Buenos Aires, Argentina; Fundación Instituto Leloir, IIBBA-CONICET, Av. Patricias Argentinas 435, C1405BWE, Buenos Aires, Argentina; Departamento de Química Biológica, FCEyN, UBA, Buenos Aires, Argentina

**Keywords:** Melanin-Tryptophan-*Pseudomonas aeruginosa*-Iron-Cystic fibrosis

## Abstract

*Pseudomonas aeruginosa* is a human opportunistic pathogen, capable of producing a wide range of metabolites, including pyomelanin. This pigment results from alterations in tyrosine catabolism. Melanin synthesis from tryptophan has never been reported in *Pseudomonas*. In this study, we describe a tryptophan-derived melanin in *P. aeruginosa* PAH, a strain that was isolated from a fibrocystic patient. PAH produced a brown pigment when grown in LB or L-tryptophan-supplemented media. Structural analysis revealed this pigment was composed by two fractions differing in NaOH solubility: a soluble one consistent of pyomelanin, and an insoluble fraction with a complex structure containing substituted indolic units. A pyomelanin inhibitor enhanced total melanin synthesis, mainly the insoluble fraction, and a tryptophan 2,3-dioxygenase inhibitor decreased pigment formation. Metabolomic profiling identified distinct indolic compounds and low levels of anthranilate in PAH cultures. Genomic and transcriptomic analyses revealed the presence of mutations and downregulation of genes related to pyoverdine biosynthesis. Furthermore, iron supplementation in the culture medium reduced melanin production. Overall, tryptophan arises as a key compound for melanin production in PAH, expanding the diversity of melanins synthesized by this genus. Furthermore, iron deprivation emerges as a critical factor triggering melanin biosynthesis, probably as a survival strategy enabling persistence in the fibrocystic lung environment.

## Introduction

Bacterial adaptability is a fundamental trait for survival in hostile environments such as the human host during infections, where multiple stressors—including oxidative molecules produced by the immune system, the presence of other microorganisms, antibiotics, and nutrient limitation can compromise cellular viability. To counter these challenges, bacteria rely on a wide range of adaptive strategies, including structural changes, metabolic shifts, and coordinated behaviors such as biofilm formation, production of antioxidant molecules, capsule synthesis, and lipopolysaccharide modification (Fang et al., 2016; Flemming et al., 2016; Whitfield et al., 2020; Needham et al., 2013).

*Pseudomonas aeruginosa* is a Gram-negative bacterium responsible for a broad spectrum of infections, particularly in immunocompromised individuals and patients with genetic conditions like cystic fibrosis (CF). Although cystic fibrosis transmembrane conductance regulator (CFTR) modulator therapies have improved clinical outcomes, certain patient groups—due to age, mutation type, or limited drug access in low- and middle-income countries—still cannot benefit from them (Taylor-Cousar et al., 2023; Guo et al., 2024).

*P. aeruginosa* expresses a variety of virulence factors, including cytotoxic pigments such as pyocyanin, tissue-degrading enzymes like elastases and collagenases, lipases, a type VI secretion system, and a strong capacity to form biofilms (Qin et al., 2022). Pigments are chemically diverse molecules with important biological functions. In the *Pseudomonas* genus, several secondary metabolites are produced, including pyoverdine (the main siderophore in *P. aeruginosa*), pyocyanin, and pyochelin. Other pigments, such as pyorubin and pyomelanin can also be found in clinical and environmental *Pseudomonas* strains.

Melanins are heterogeneous polymeric pigments formed mainly by oxidative polymerization of indolic or phenolic compounds. The main types include DOPA-melanin (also called eumelanin) and homogentisate-derived pyomelanin. Tyrosine-derived intermediates are common biosynthetic precursors for both DOPA-melanin and pyomelanin, although other substrates such as malonyl-CoA may also contribute to melanin biosynthesis (Pavan et al., 2020; Lorquin et al., 2022).

In addition, other forms of melanin, such as allomelanin, can be synthesized by various microorganisms. Allomelanin is derived from the oxidative polymerization of nitrogen-free precursors, including 4-dihydroxyphenylacetic acid, catechol, dihydroxynaphthalene (DHN), protocatechualdehyde, or tetrahydroxynaphthalene, and has been identified in both bacterial and fungal species (Cao et al., 2021; Martinez et al., 2023). For instance, within the *Pseudomonadaceae* family, catechol-based melanin production has been reported in certain *Azotobacter chroococcum* strains under nitrogen-fixing conditions in the presence of oxygen (Shivprasad and Page, 1989; Herter et al., 2011).

These observations underscore the metabolic versatility of bacteria in synthesizing melanin through distinct pathways and from diverse chemical precursors. Notably, a recent study demonstrated that tryptophan can also serve as a physiological substrate for melanin biosynthesis in the photosynthetic bacterium *Rubrivivax benzoatilyticus* JA2, resulting in a pigment termed tryptophan-derived melanin (Ahmad et al., 2020).

Here, we describe a novel melanin biosynthetic route in a clinical *P. aeruginosa* isolate (PAH) recovered from a child diagnosed with CF, in which tryptophan serves as a precursor for melanin production, mainly via anthranilate and indolic compounds, under iron-depleted conditions. This finding expands the current understanding of pigment diversity and metabolic plasticity in *P. aeruginosa* and the role of iron availability in melanin production.

## Material and methods

### Bacterial strains and isolates

*P. aeruginosa* PAH was isolated from a pediatric patient with cystic fibrosis during routine controls in the General Children‘s Hospital Dr. Pedro Elizalde under the Ethical Committee Approval number PRISBA 4466, with informed consent of participant or his/her legal guardians. Isolates were identified by routine culture followed by MALDI-TOF identification. Bacterial protein extraction for MALDI-TOF MS was performed as previously described for microbial identification. Briefly, bacterial cells grown aerobically for 16–20 h on Mueller-Hinton agar were resuspended in 70% ethanol and mechanically disrupted using 0.1 mm glass beads. Cell lysates were treated with a 1:1 mixture of 70% formic acid and 100% acetonitrile. An aliquot of the supernatant was applied to the MALDI target plate, overlaid with a matrix solution containing α-cyano-4-hydroxycinnamic acid (CHCA), and air-dried. Spectra were acquired using the VITEK MS platform (bioMérieux), based on a Shimadzu Biotech Axima Assurance mass spectrometer (software version 2.9.5.6), under default acquisition and calibration settings provided by the manufacturer.

The reference *P. aeruginosa* PAO1 strain, a wound isolate (Stover et al., 2000), and PAO1 *hmgA*\*, an isogenic pyomelanin-producing mutant constructed by CRISPR-Cas technology by generating a premature stop codon in the *hmgA* gene of P. aeruginosa PAO1 (Diaz Appella et al., 2024), were used for comparisons.

### Bacterial culture conditions

Routine growth of *P. aeruginosa* was performed using triplicate overnight pre-cultures in LB medium under aerobic conditions (1:10 culture to Erlenmeyer flask volume and 150 rpm agitation) at 37 °C. To perform the growth curves, after centrifugation and washing with PBS, cells were resuspended in the same medium and used to inoculate cultures to an initial OD_600nm_ of 0.025. For oxidative stress resistance and HPLC analysis, succinate minimal medium supplemented with 2 mM tryptophan was used. Such medium consisted of 0.5% w/v Na_2_HPO_4_, 0.3% w/v KH_2_PO_4_, 0.15% w/v (NH_4_)_2_SO_4_, 0.6% w/v sodium succinate, 0.2% w/v glycerol and 0.3% w/v MgSO_4_, adjusted to pH 7.2.

### Transmission Electron Microscopy analysis

The cell shape and the pigment of PAO1, *hmgA\**, and PAH strains grown for 24h in LB medium were analyzed by transmission electron microscopy (TEM). Cells were harvested, washed twice in PBS, and fixed in 2.5% (w/v) glutaraldehyde. Bacterial samples or purified pigments were negatively stained with 2% aqueous uranyl acetate. Suspensions were placed onto 200-mesh copper grids coated with an LR White membrane and examined using a Zeiss EM109T transmission electron microscope operated at 80 kV. Images were captured with a Gatan ES1000W camera at the Institute of Cell Biology and Neurosciences ‗Professor Eduardo de Robertis‘, School of Medicine, University of Buenos Aires.

### Cell culture conditions

The A549 lung epithelial cell line was maintained in Dulbecco‘s Modified Eagle‘s Medium (DMEM), supplemented with 10% heat-inactivated fetal bovine serum, 100 U/mL penicillin, and 100 µg/mL streptomycin (Sigma). Cells were incubated in 25-cm^2^ tissue culture flasks at 37 °C under a humidified atmosphere containing 5% CO_2_.

### Genome sequencing and analysis

Genomic DNA from *P. aeruginosa* PAH was obtained using a commercial kit (GenElute™ Bacterial DNA Kit, Sigma-Aldrich). The purified DNA was sequenced using an Illumina NovaSeq 600 system (Novogen, California, USA). The assembly was performed using *SPAdes* genome assembler v3.15.5 (Prjibelski et al., 2020). The genome was deposited in the National Center for Biotechnology Information (NCBI) database under the accession number JBOEHO000000000. For SNPs analysis, the PAO1 genome (GCA_000006765.1) was used as reference. Thus, a VCF file was generated using *BCFtools*, and the variants and their effects were annotated employing the *snpEff* tool (Cingolani et al., 2012). To compare the genomic distance, the genomes of several reference strains were downloaded from NCBI (GenBank accessions: PAO1: GCF_000006765.1; PA14: GCA_045689255.1; PAK: GCA_000408865.1; NCGM2: GCA_000284555.1; PACS2: GCA_000168335.1; SG17M: GCA_020978345.1; RP73: GCA_000414035.1; DK2: GCA_000271365.1; LESB58: GCA_000026645.1; and *P. paraeruginosa* PA7: GCA_000017205.1). The alignment for subsequent phylogenetic inference was generated with Parsnp v2.0.3, using PAO1 as reference. From the Parsnp binary file, the complete core-genome alignment was exported with HarvestTools. A maximum-likelihood tree was inferred with IQ-TREE 2, and the resulting tree was rooted at PAO1 in FigTree v1.4.4.

### Pigment isolation, quantification, and inhibition analysis

Total melanin pigment was purified as described previously (Diaz Appella et al., 2024). Briefly, 50 mL of 24 h cell-free culture supernatants were acidified (pH 2) using 5 M HCl. After 24 h incubation at room temperature in the dark, the precipitated melanin was centrifuged, washed twice with 20 mL of ultrapure (MilliQ) quality water and once with acetone:ethanol solution (1:1), and resuspended in water at 100 °C for 15 min. After centrifugation, the precipitate was washed with the same volume of water and freeze-dried to get total purified melanin. To obtain the insoluble melanin (approximately 53% of the initial dry weight, depending on the culture medium and supplements), two washes with 20 mL of NaOH 0.5 M and three washes with distilled water were performed. The insoluble fraction of the pigment was re-lyophilized. To evaluate melanin production using different compounds as precursors, cultures were grown for 24 h in minimal medium supplemented with varying equimolar proportions of potential biosynthetic precursors, including L-tyrosine, L-tryptophan, L-phenylalanine, indole-3-acetic acid, anthranilic acid (Sigma-Aldrich). In all cases, the total concentration of supplemented compounds was adjusted to 2 mM, regardless of the combination used. Iron sulfate and three melanin synthesis inhibitors were tested: sodium azide (Sigma-Aldrich), kojic acid (Sigma-Aldrich), bicyclopyrone (ACURON™ UNO; Syngenta Agro S.A.), and 6-fluoro-3-[(1E)-2-(3-pyridinyl)ethenyl]-1H-indole (Sigma-Aldrich), an inhibitor of tryptophan 2,3-dioxygenase (TDO). To ensure that the selected concentrations did not affect cell viability, growth curves were performed for sodium azide and kojic acid across a range of concentrations (0.1-10 mM and 0.3-5 mM, respectively). Bicyclopyrone was used at 0.1 mM, a concentration previously reported (Diaz Appella et al., 2024). For the TDO inhibitor, two concentrations were tested (0.005 and 0.05 mM), as neither impacted viability, and the highest was selected for further assays. All melanin production assays were conducted in quadruplicates using 25 mL LB cultures. Pyocyanin production was evaluated using the standard chloroform-based extraction protocol (Kurachi 1958). Each assay was conducted in triplicate.

### Supernatant HPLC and nuclear magnetic resonance (NMR) metabolomic analysis

*P. aeruginosa* strains were grown in succinate minimal medium (hereinafter, MM), either without supplementation or supplemented as specified, at 37 °C. Samples were taken at 18 h of growth, and centrifuged for 10 min at 5,000 g and 4 °C. The supernatants were filtered through a 0.2 μm pore size sterile syringe filter (Millipore, Germany). HPLC separation of the supernatants was carried out on an Agilent 1200 HPLC with a diode array detector using a Poroshell 120 EC C18, 2.7 µm, 4.6 x 150 mm column (Agilent, Germany) fitted with a guard column. A linear gradient from 0.1% phosphoric acid in water (solution A) to acetonitrile (solvent B) for 30 min with an additional 5 min of 100% solvent B at a flow rate of 1 mL/min was used, and 10 µL were injected. Peaks were detected at 210, 230, 260, 280, and 310 nm wavelengths.

For supernatant targeted NMR profiling analysis, each sample (20 mg) was reconstituted in 0.5 mL of sodium phosphate buffer prepared in D_2_O (pH 7.40 ± 0.05), containing 3-trimethylsilyl-[2,2,3,3-^2^H_4_]-propionate (TSP) at a final concentration of 0.33 mM, and then transferred to 5 mm NMR tubes. All NMR measurements were carried out at 298 K on a Bruker Avance III spectrometer equipped with a proton resonance frequency of 600.1 MHz (Bruker BioSpin GmbH & Co. KG, Ettlingen, Germany). The experimental protocols as well as data processing and analysis were performed in accordance with previously established methodologies (Pellizza et al., 2022; Avalos et al., 2025). For metabolite quantification, only well-resolved, non-overlapping peaks of the identified metabolites were integrated. ^1^H-NMR 1D spectra were acquired using a standard Bruker 1D NOESY pulse program with pre-saturation during relaxation delay and mixing time, and spoil gradients (noesygppr1d). The NMR spectral areas (AUC) of the assigned metabolites were normalized using probabilistic quotient normalization (PQN). Statistical significance was evaluated by the Wilcoxon rank-sum test, taking p<0.05 as significant. Metabolite assignment was performed using the freely accessible electronic databases: HMDB (Human Metabolome Database) (Wishart et al., 2022) and BMRB (Biological Magnetic Resonance Data Bank) (Hoch et al., 2023). Assignment was subsequently confirmed through two-dimensional (2D) NMR spectra, including heteronuclear single-quantum coherence (HSQC) and total correlation spectroscopy (TOCSY), as well as by comparison with authentic reference compounds anthranilate and indolacetate (Sigma).

### Structural analysis of the pigment

Fourier-transform infrared (FT-IR) spectroscopy with attenuated total reflectance (ATR) was conducted using a Nicolet IS50 spectrometer (Thermo Scientific, Waltham, MA, USA). For UV–Vis spectrophotometric analysis, purified melanin was initially dissolved at 1 mg·mL⁻ ¹ in 0.2 M NaOH, then diluted to 0.03 mg·mL⁻ ¹ prior to measurement. NMR analysis was performed on a Bruker Avance Neo 500 MHz spectrometer (Bruker BioSpin GmbH & Co. KG, Ettlingen, Germany) in solution samples (soluble pigment fractions) using standard NMR analysis and on DMSO swollen pigment samples employing a HR-MAS probe. For solution NMR, 15 mg of melanin was dissolved in 0.6 mL of 0.05 M NaOH in D_2_O. ^1^H and ^13^C (1D and 2D) NMR spectra were recorded using the residual solvent signal as an internal reference (^1^H: 500 MHz, ^13^C: 125.4 MHz), with the ^1^H spectrum acquired using the pulse sequence zg30 and the ^13^C spectrum using the pulse sequence zgpg45. The 2D spectra acquired were a Pure-Shift HSQC spectrum (kp_rtPS_gHSQC_ts4) and a COSY spectrum (cosygpqg60). For HR-MAS analysis, about 15 mg of pigment was swollen with 50 μL of DMSO into a 4 mm zirconia rotor. Additional DMSO was added as needed to fill the rotor, and excess solvent was removed with paper tissue before closure of the rotor with a Kel-F cap. Spectra were acquired on a triple channel HCND HR-MAS probe with rotors spun at 4000 Hz at the magic angle axis and at 298 K, using the residual solvent signal as an internal reference. ^1^H spectra were acquired using a CPMG pulse program (cpmgpr1d), while 2D spectra included COSY (cosygpppqf) and phase-edited HSQC (hsqcedetgpsp.3) spectra.

### RNAseq data analysis

Total RNA extraction of PAH and PAO1 LB cultures incubated for 18 h at 37 °C was performed using the Trizol method followed by a commercial kit (Total RNA extraction kit—RBC bioscience). Samples were treated with DNase I (Promega). To improve the quality of the readings, ribosomal RNA was depleted from the samples by Novogen Services (CA, USA), followed by library construction. Mass sequencing was performed using the Illumina Novaseq 6000 platform with a paired-end protocol (Novogen Services; CA, USA). Independent RNA extraction and libraries replicates (PAH N=4; PAO1 N=3) were used for each strain. Reads were preprocessed by Novogen company, including the adapter trimming. Read quality was evaluated using the FastQC tool. Reads alignment and assembly, transcript identification, and abundance quantification were carried out using the Rockhopper software using the uploaded *P. aeruginosa* PAO1 (AE004091.2) closed genome (Tjaden et al., 2020). Reads were normalized per kilobase per million mapped reads. To verify concordance between the independent replicates, a Spearman correlation analysis of the normalized counts was performed. Genes were only considered as differentially expressed with *P* < 0.05 and *Q* < 0.05 and a Fold Change (FC) ≤ -1.5 and ≥ 1.5. *q*-values are false discovery rate (FDR) adjusted *p*-values, using the Benjamini–Hochberg method. *P* < 0.05 and *Q* ≤ 0.15 and ≥ 0.05 were considered indicative of a trend toward statistical significance. Genes were sorted into functional classes using the KEGG (Ogata et al., 1999), MetaCyc (Caspi et al., 2016), and String (Szklarczyk et al., 2019) tools. For quantification and classification, tRNA transcripts were not considered. RNAseq data can be found in the National Center for Biotechnology Information (NCBI) database under the accession number PRJNA1268010.

### Oxidative stress resistance analysis

To assess *P. aeruginosa* PAH resistance to oxidative stress, growth inhibition assays were performed on MM agar (with or without 2 mM tryptophan supplementation) and LB agar. Aliquots of 100 µL from overnight cultures adjusted to an OD_600nm_ = 1 were plated in quadruplicate. Sterile filter paper discs (6 mm diameter) were placed onto the agar surface and soaked with 5 µL of 30% H_2_O_2_ or 27.6% paraquat ( 1,1’-dimethyl-4,4’-bipyridilium dichloride, *Sigma Agro*). Once the plates had dried, they were incubated overnight at 37 °C and inhibition zones were measured to quantify the oxidative stress resistance.

### Melanin cytotoxicity

The purified melanin lethal dose concentration 50% (LD_50_) was determined using the colorimetric, 3-(4,5-dimethylthiazol-2-yl)-2,5-diphenyltetrazolium bromide (MTT, Sigma) assay. Briefly, A549 cells (a human adenocarcinoma epithelial cell line) were cultured at 37 °C under 5% CO2 in Dulbecco‘s modified eagle medium (DMEM, Gibco) supplemented with 1% penicillin–streptomycin (Pen-Strep, Gibco) and 10% fetal bovine serum (FBS, Gibco). 96-multiwell plates containing 90% confluent monolayer of A549 were incubated for 2 h with DMEM supplemented with 10% (v/v) FBS and serial dilutions of a stock solution of melanin (10 mg/mL). Then, each well was washed twice with PBS, and the A549 cells were cultured in fresh medium containing 0.5 mg/mL MTT for a further 4 h period. The blue formazan products in the A549 cells were dissolved in DMSO and spectrophotometrically measured at a wavelength of 550 and 690 nm. All experiments were replicated in quadruplicates.

### Cytokine production

A549 monolayers were cocultured with bacterial supernatant or resuspended melanin, and their cytokine production was analyzed using specific enzyme-linked immunosorbent assay (ELISA). Briefly, 96-multiwell plates containing cells at 90% confluent growth in similar conditions to those used for the cytotoxicity assay were centrifuged and washed with PBS. Then, 200 µL of DMEM supplemented with 10% FBS and the appropriate concentration of melanin were added to the cells, which were then cultured (37 °C, 5% CO_2_) for 16 h. After incubation, supernatants were collected and stored for one day at -20 °C, to evaluate the cytokine concentration using specific commercial ELISA kits (Sigma Aldrich).

### Statistical analysis

For multiple comparisons, the significance of the differences between means was evaluated through non-parametric one-way analysis of variance (ANOVA) with a Kruskal-Wallis and Dunn’s multiple comparisons test, and unpaired and nonparametric Mann–Whitney correction, with confidence levels at >95% (i.e., *P*<0.05 was considered significant). For two-mean comparisons, *t*-test analysis with the corresponding correction was performed, with confidence levels at >95%. All the analyses were performed using GRAPH-PAD PRISM version 8.0.2 (Dotmatics, Boston, MA, USA) for Windows.

## Results

### The *P. aeruginosa* PAH isolate produces a brownish pigment from tryptophan

*P. aeruginosa* PAH was obtained from a pediatric patient with CF in the context of a coinfection with *Staphylococcus aureus.* The infection in this patient was considered chronic according to the Argentinian Society of Pediatrics’ criteria. PAH was isolated in the General Pediatric Hospital ―Dr. Pedro Elizalde‖ (Buenos Aires, Argentina) and identified as a *P. aeruginosa* strain by routine culture and MALDI-TOF analysis (Fig. S1). PAH presented an unusual color in LB medium, different from PAO1 and the pyomelanin producer PAO1 *hmgA\** mutant strain, but showed a similar growth compared to these strains after 24 h (Fig. 1A and B).

**Figure 1.**
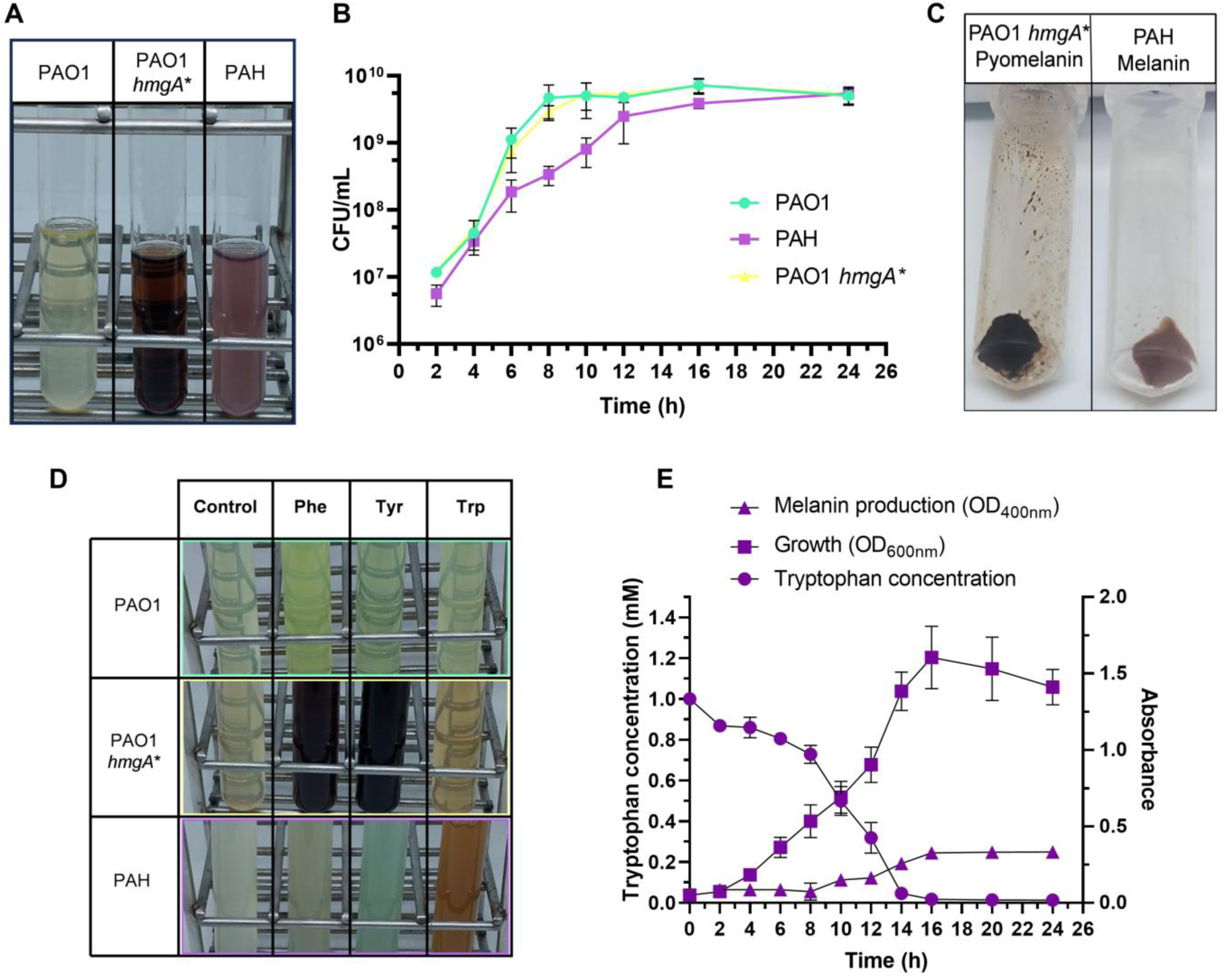
Growth and melanin production in the PAH clinical isolate. (A) Shown is the supernatant of *P. aeruginosa* strains grown for 24 hours in LB. (B) Growth curves of PAH, PAO1 and PAO1 *hmgA*\* in LB medium. (C) Purified pyomelanin from PAO1 *hmgA*\* and melanin from PAH. Both strains were grown in LB medium. (D) Shown is the supernatant of *P. aeruginosa* strains grown for 24 h in succinate minimal medium (MM) supplemented with 2 mM of phenylalanine (Phe), tyrosine (Tyr) or tryptophan (Trp). (E) Interplay between growth of PAH in minimal medium (MM) supplemented with 2 mM tryptophan, and melanin and tryptophan concentrations. For all panels, each assay was repeated in triplicate.

To determine whether the color of the PAH strain was due to a new pigment or a combination of different ones, we performed different organic extractions of the supernatant of PAH and PAO1 cultures in LB. These analyses showed similar pyocyanin production between PAH and PAO1 (A_520nm_/OD_600nm_ 0.039 and 0.030 for PAH and PAO1, respectively), but also the presence of brownish pigments solely in the PAH supernatant. Thus, acid precipitation of the PAH supernatant was performed, resulting in the purification of a brownish pigment (Fig. 1C). Although pyomelanin is typically soluble in NaOH solution, the PAH pigment powder separated into two fractions in NaOH solution, a soluble and an unexpected insoluble fraction. To understand the biosynthetic origin of the brown pigment, we cultured PAH in minimal medium supplemented with individual aromatic amino acids - tyrosine, phenylalanine and tryptophan, observing that the pigment was only produced in the presence of tryptophan (Fig. 1D). Time course analysis revealed that in minimal medium supplemented with tryptophan, the pigment was detectable at approximately 10 h post-inoculation, increased steadily until 16 h, and then reached a *plateau*, remaining stable up to 24 h (Fig.1E). At 24 h, the total pigment was purified and quantified, yielding a concentration of 0.15 mg/mL ± 0.03 mg/mL. Importantly, pigment production was coincident with tryptophan consumption (Fig. 1E).

We determined total melanin production in the presence of other tryptophan metabolism-related compounds: anthranilate, and indole acetic acid alone or combined with tyrosine or tryptophan, along with other control amino acids including lysine and aspartic acid (Fig. 2A). Supplementation of both anthranilic acid and tryptophan resulted in similar melanin production (0.10 mg/mL ± 0.01) to tryptophan supplementation alone, while no or low production was detected for the other combinations (Fig. 2A). Interestingly, the combination of tryptophan and tyrosine resulted in the production of 0.09 mg/mL ± 0.01 of total melanin, while other combinations did not show relevant pigment production in *P. aeruginosa* PAH (Fig. 2A). Upon evaluating the proportion of soluble versus insoluble fractions under each condition (Fig. 2B), we observed that supplementation with tryptophan led to almost exclusively insoluble fraction production (90%, hereinafter trp-mel), whereas the addition of tyrosine increased the proportion of the soluble fraction (67%, hereinafter pyo-mel). Additionally, we analyzed the impact of various inhibitors targeting melanin-associated enzymes to gain insights into the potential biosynthetic pathways involved. The addition of sodium azide and kojic acid, which prevent eumelanin synthesis through inhibition of laccase and tyrosinase activity, respectively, had no observable effect on total pigment or trp-mel production. In contrast, treatment with bicyclopyrone, an inhibitor of the 4-hydroxyphenylpyruvate dioxygenase involved in pyomelanin synthesis, led to an increase in total melanin production with an increment in trp-mel proportion. Finally, inhibition of tryptophan 2,3-dioxygenase (TDO), encoded by *kynA*, resulted in a decrease in total pigment synthesis and in the trp-mel fraction, suggesting the involvement of tryptophan-derived intermediates in the biosynthetic process (Fig. 2C). We also performed transmission electron microscopy (TEM) analysis to investigate the morphology of PAO1, *hmgA\**, and PAH strains when cultured in LB medium. Cells of all strains exhibited similar bacillar morphology. In melanin-producing strains, *hmgA\** and PAH, the pigment could be visualized associated with the cells. In *hmgA\**, pyomelanin showed a laxer structure compared to PAH melanin, which presented a compact appearance (Fig. 2D). TEM analysis of the melanin powder after acid precipìtation from the supernatant of cultures in LB showed a more compact, packing structure in the total melanin of PAH than in the pyomelanin of PAO1 *hmgA*\* (Fig. 2E), providing preliminary evidence of the differences between the polymers.

**Figure 2.**
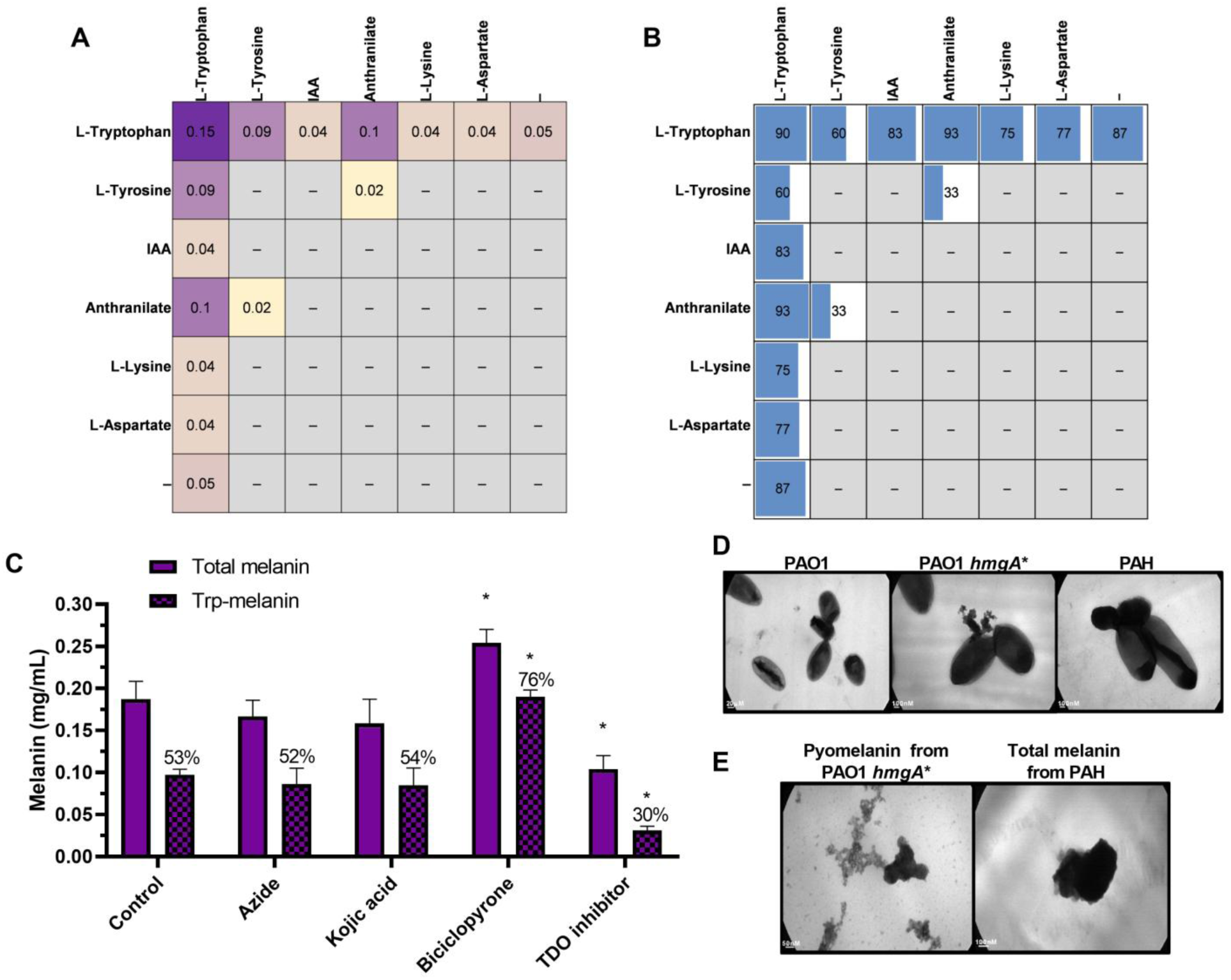
Substrates and metabolic pathways involved in PAH melanin production. (A) Matrix of total melanin production (mg/mL) after 24 h growth of PAH in minimal medium supplemented with 1 mM of the compound indicated at the top of each column and 1 mM of the compound indicated at the left of each row.‖IAA‖ stands for indole-3-acetic acid; ―–‖ indicates no supplementation and was used as the control condition. (B) Matrix of trp-mel production (%) relative to the total melanin produced under each condition shown in panel A. (C) Total melanin and Trp-mel production after 24 h growth of PAH in LB medium in the presence of canonical melanin synthesis inhibitors: sodium azide and kojic acid (DOPA-melanin inhibitors), bicyclopirone (a pyomelanin synthesis inhibitor), and a tryptophan 2,3-dioxygenase inhibitor (TDO inhibitor). Bars indicate total melanin and Trp-melanin levels, and the percentages correspond to Trp-melanin relative to total melanin for each condition. In the ―Control‖ condition no inhibitor was added. Significant differences (*: P<0.05) are shown compared to the control condition. (D) Transmission electron microscopy images of the indicated strains cultured in LB medium. (E) Transmission electron microscopy images of the purified pigments extracted from LB cultures of each strain. Scale bars are indicated in each panel. For all panels, each experiment was performed in quadruplicate.

### Targeted NMR profiling reveals strain-specific tryptophan catabolic signatures

To further characterize the observed pigment, we analyzed intermediates present in the culture supernatant of PAH and PAO1 in minimal medium supplemented with tryptophan by HPLC. Higher consumption of the tryptophan was detected in PAH than in PAO1. Based on absorbance spectra, distinct compounds were detected in PAH that were not observed in PAO1 (Fig. S2). These spectra were compatible with indole species. Notably, these compounds were not detected when PAH was cultured in media supplemented with tyrosine or phenylalanine (Fig. S3 and S4), indicating that their accumulation is specific to tryptophan supplementation.

Next, we performed targeted NMR-based metabolomics to investigate tryptophan-derived metabolites of PAO1 and PAH cultured in minimal medium supplemented with tryptophan. This analysis enabled tryptophan and anthranilic acid to be confidently assigned, with anthranilic acid dominating the NMR spectrum of the PAO1 strain and PAH exhibiting a significant decrease in extracellular anthranilate levels (Fig. 3 and Fig. S5). Additionally, indole derivatives were detected in the PAH strain, with at least two potential different indolic compounds that were absent in PAO1, as assessed by heteronuclear single quantum coherence (HSQC) spectroscopy (Fig. 3, Fig. S5). The observed pattern indicates a potential metabolic rerouting of tryptophan catabolism in the PAH strain, which may favor the accumulation of indole-derived compounds.

**Figure 3.**
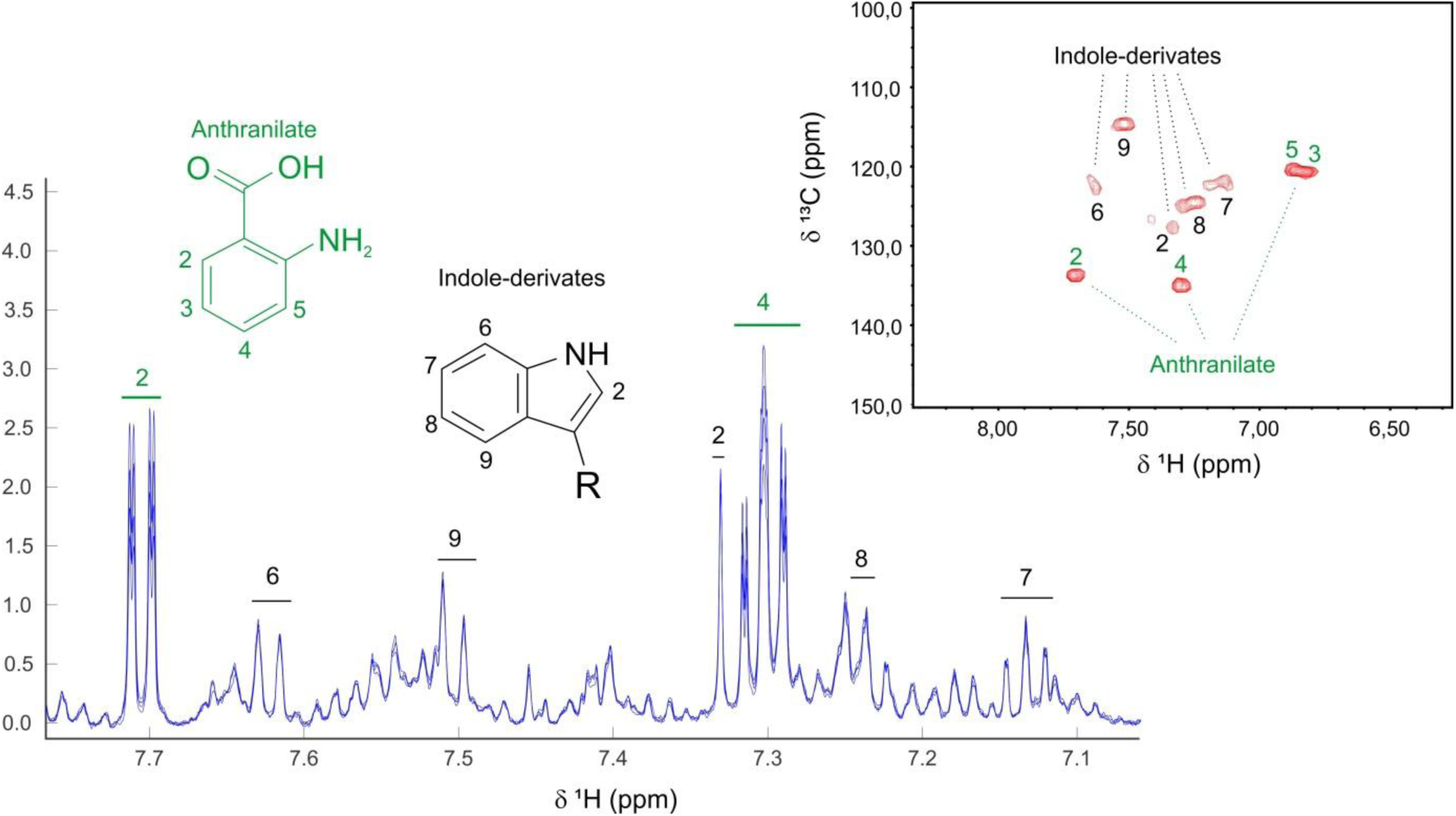
NMR assignments of anthranilate and indole-derived metabolites identified in supernatants of the H strain. Overlaid 1 H-NMR spectra of four replicates showing resonances assigned to anthranilate (green) and indole-derived compounds (black). Proton numbering corresponds to the annotated molecular structures. The 2D 1 H– 13 C HSQC (inset) further confirms the assignments, displaying characteristic cross-peaks for anthranilate and indole-related metabolites.

### PAH melanins present different structural features

Given the uncommon nature of our finding, we carried out a structural analysis of trp-mel and pyo-mel compared to pyomelanin from PAO1 *hmgA*\*. As described above, trp-mel and pyo-mel showed different NaOH solubility, therefore, we analyzed the different fractions to understand the structural identity of the pigments. The FTIR (ATR) spectrum of total melanin (performed in solid state) showed absorptions at 3267 (O–H from carboxylic acid, phenol, alcohol), 2924 (C–H from aromatic moieties), 2854 (C–H associated to sp^3^ C), 1631 (aromatic C=C overlapped with C=O from carboxylate and C=O from quinone), 1530 (aromatic C=C), 1454 (O–H), 1214 (C–O in phenol or phenol ether), and 1040 cm^-1^ (C–O in primary alcohol or ether) (Fig. 4A). Some of these signals were coincident with those found in pyomelanin from different strains of *P. aeruginosa*, including PAO1 *hmgA*\* (Diaz Appella et al., 2024). However, some differences in terms of intensity were observed, since more intense absorptions were detected for aromatic C-H (2924 cm^-1^) and aliphatic C-O bonds (1040 cm^-1^) in comparison with PAO1 pyomelanin. Moreover, the intensity ratio corresponding to phenol: primary aliphatic C-O bands indicated an increase in the aliphatic ether content. The presence of absorption bands related to nitrogen-containing functionalities could not be resolved from oxygen-related bonds at 3267 and 1530 cm⁻¹.

**Figure 4.**
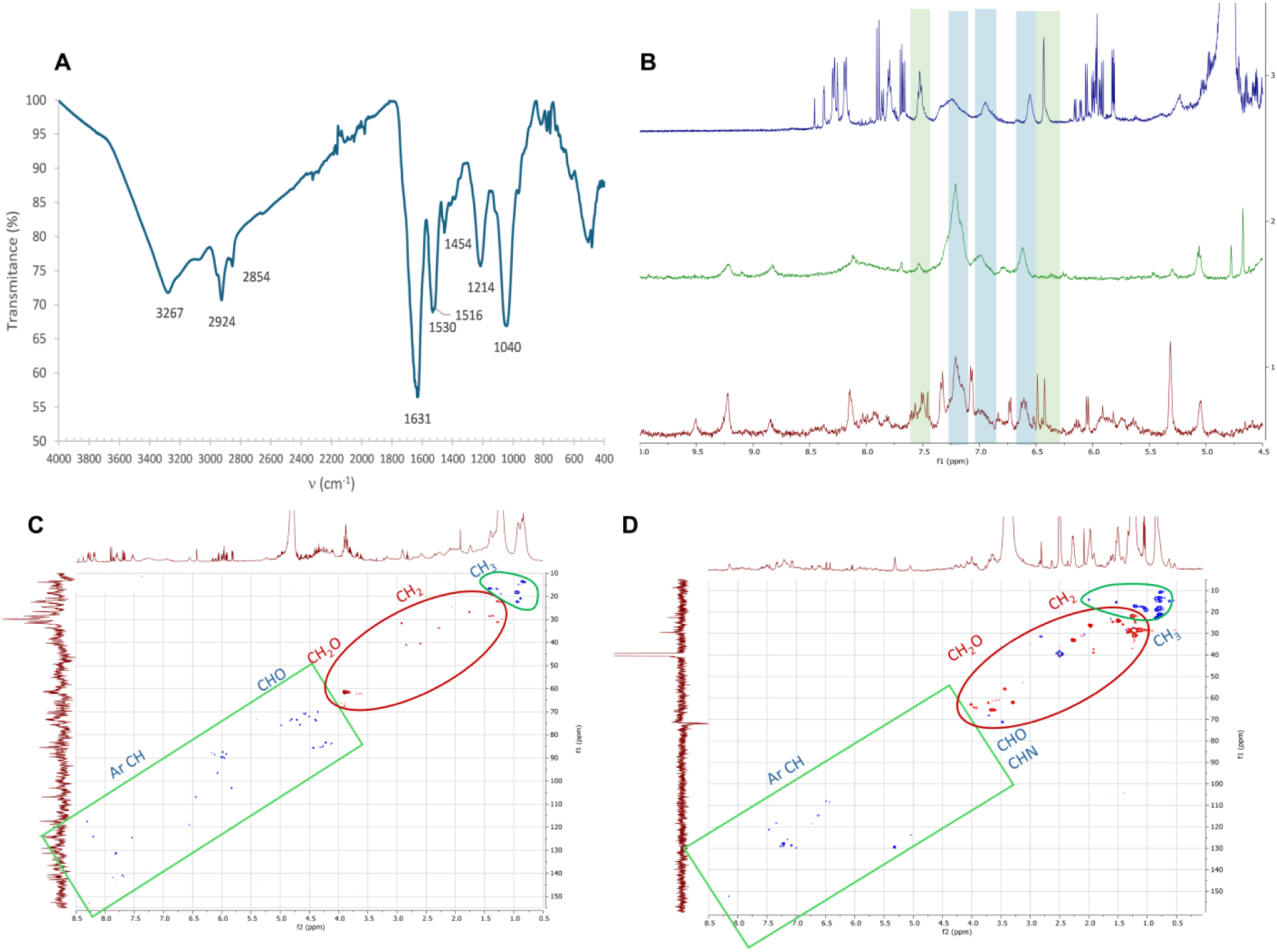
(A) FTIR (ATR) spectrum of PAH pigment; (B) ^1^H NMR spectra of pyo-mel (D_2_O, top), PAO (HR MAS – CPMG, DMSO-*d*_6_, middle), Trp-mel (HR MAS – CPMG DMSO-*d*_6_, bottom); (C) HSQC spectrum of pyo-mel (D_2_O), (D) HSQC spectrum of trp-mel pigment (DMSO-*d*_6_)

Next, different NMR techniques were performed on both PAH melanin fractions. The ^1^H NMR spectrum of the pyo-mel from PAH showed a complex overlapping of broad and sharp resonances (Fig. 4B). The aromatic region exhibited a set of three broad signals consistent with pyomelanin, where the signal at 6.55 ppm was attributed to benzoquinone moieties, and the resonances at 6.95 and 7.15-7.35 ppm were assigned to aromatic protons in rings linked through C–C and C-O-C phenol ether bonds, respectively (Diaz Appella et al., 2024). In addition, a broad singlet at 6.43 ppm, with a cross peak at 106.9 ppm in the ^1^H-^13^C HSQC spectrum (Fig. 4C), could be attributed to disubstituted catechol or resorcinol (1,2-dihydroxyphenyl or 1,3-dihydroxyphenyl) moieties included in the polymeric structure (Ryl et al., 2019). This fact was in agreement with the oxidative condensation of catechol-like residues in aqueous solution, leading to C-C and ether C-O-C linkages between aromatic rings and a ring opening reaction promoted at alkaline pH, yielding aliphatic chains and carboxylic acids (Sanchez Cortés et al., 2001). Aliphatic signals in the range 4.0 – 5.0 ppm indicated the presence of CHO groups that were close to CH_2_OH residues at 3.5 – 4.0 ppm, as indicated by the COSY ^1^H ^1^H spectrum. Additional methylene groups were recorded in the range 1.2 – 3.0 ppm, while methyl groups were observed at 1.4 – 0.8 ppm. The ^13^C NMR spectrum of the pyo-mel showed relevant signals, namely carbonyl resonances from carboxylic acids and esters (172 – 173 ppm), quinone or quinolone units (181 ppm), quaternary C–O (weak signals at 145-160 ppm), and C-H from phenol residues (102-140 ppm). At a higher field, CHO and CH_2_OH groups were recorded at 61.0-89.5 ppm, while CH_2_ and CH_3_ groups were detected at 22-41 and 13-22 ppm, respectively. The above assignments for the aqueous, soluble polymeric structure agreed with the solubility behaviour of a melanin containing carboxylic acids and phenols that would be deprotonated and solubilized in alkaline conditions, with aromatic resonances in the NMR spectra shifted to higher fields. Additionally, with the assistance of 2D NMR experiments, a set of five sharp aromatic resonances were assigned to 2-heptyl-4(1H)-quinolone (HHQ), a quorum-sensing signal of *P. aeruginosa* (Camilios-Neto et al., 2024).

To better characterize the polymeric structure, HR-MAS NMR experiments (1D and 2D) were carried out on the trp-mel from PAH and pyomelanin from PAO1 *hmgA\** (Fig. 4D). The spectra showed some common features between reference pyomelanin and the trp-mel, suggesting common polymeric domains. However, a more complex pattern was detected for the insoluble PAH pigment trp-mel. In the aromatic region, some additional indolic moieties with different oxidation degrees could be associated to resonances previously reported for polydopamine (8.14, 7.08 ppm) (Liebscher et al., 2013), 5,6-dihydroxyindole (6.73 ppm) and its carboxylic acid derivative (7.21 ppm) present in melanin (Restaino et al., 2022). One pair of distinctive signals at 6.04 and 6.73 ppm exhibited a cross peak in the COSY spectrum that could suggest an indolic unit with unsubstituted C-2 and C-3 positions. Another intense correlation signal in the HSQC spectrum was detected at 5.31 (broad singlet) and 129.4 ppm, which could originate from a conjugated vinylic unit, but could not be unequivocally assigned.

In summary, the PAH pigment seems to be composed of a complex mixture including a soluble pyo-mel fraction structurally similar to the PAO1 pyomelanin, together with some additional units of hydroxy-substituted aromatic moieties. The insoluble trp-mel main fraction had a more heterogeneous composition that probably incorporated a new and complex array of indolic units substituted with hydroxyl and carboxylic acid groups.

### Iron deprivation is a key factor for PAH trp-mel production

PAH‘s genome was sequenced to understand which metabolic branches are involved in PAH pigment production. Whole genome analysis revealed that the PAH genome has around 6.37 Mbp with a mean G+C content of 66%., and that PAH belongs to the PAO1 group, having the shortest genomic distance to the LESB58 strain (Fig. S6).

The predicted coding sequences for the enzymes involved in the aromatic amino acid pathways showed no differences between PAH and PAO1 (Table S1). All genes encoding tryptophan related enzymes were conserved, including the anthranilate, catechol, and indole branches (*kynA, kynB, kynU, antABC, amiEBCRS* operon) (Table S1). No extra tyrosinase, laccase, or peroxidase were observed. However, upon analysis of other key metabolic pathways, we identified multiple mutations in PAH genes associated with pyoverdine biosynthesis and corresponding iron acquisition. Mutations included insertions, deletions causing frameshifts, and premature stop codons (Fig. 5A). Since iron is a key nutrient and pyoverdin acts as the main siderophore in *Pseudomonas* species, we hypothesized that, due to these mutations, the PAH strain was under iron starvation despite the presence of iron in the culture medium. PAH pigment production was tested again in LB (which is estimated to contain around 5.8 µM iron according to Yan et al., 2013) supplemented with extra FeSO_4_ in the range of 1 up to 100 µM. Interestingly, under these conditions PAH pigment production diminished in an iron dose-dependent manner (Fig. 5B).

**Figure 5.**
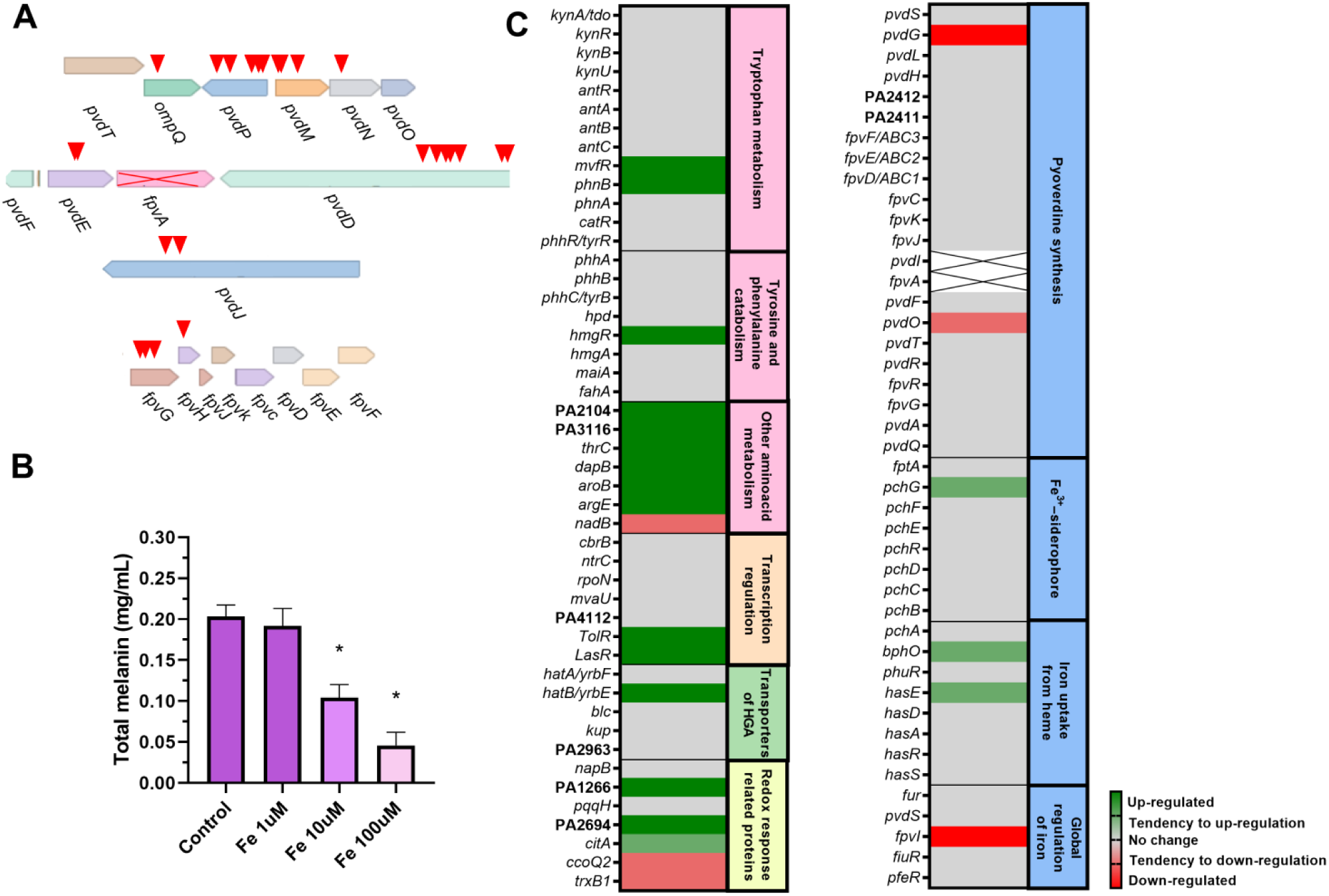
Pyoverdin (*pvd*) cluster modifications and their effects on melanin production and gene expression. (A) Gene organization of the *pvd* cluster. Red arrows indicate regions carrying mutations that cause a frameshift, and crosses indicate complete gene deletions. (B) Total melanin production in minimal medium supplemented with increasing concentrations of FeSO. Data represent the mean of four independent replicates; significant differences (*: P<0.05) are indicated relative to the control condition. (C) Heatmap of transcriptomic profiles showing relative expression levels of selected genes in PAH compared to PAO1 grown in LB medium during 20 hours. Crossed-out indicates no detected expression in PAH.

Transcriptomic analysis showed pyoverdin cluster genes *pvdP, pvdO,* and *pvdG* were downregulated while the pyochelin-related genes *pchG* and *bphO* were upregulated (Fig. 5C). Moreover, no expression was detected for *pvdE*, *pvdD*, *pvdJ*, and also for two genes related to iron transport through pyoverdin, *fpvA* was even absent in PAH‘s genome (Fig. 5C).

Although no differences in the expression *hpd* and *hmgA* related to tyrosine catabolism were detected, the *hmgR* and *hatB* coding genes were upregulated in PAH (Fig. 5C).

In addition, several genes related to redox homeostasis were found to be altered in PAH. Genes coding for a putative thioredoxin, an oxidoreductase (homologous to PA2694 and PA1266, respectively), and cytochrome *citA* were upregulated; while *ccoQ2* and *trxB1* were repressed (Fig. 5C). Several genes related to amino acid metabolism were differentially expressed in PAH, among them *aroB*, *argE* and *thrC* were upregulated while *nadB* and *cysE* were downregulated (Fig. 5C). Interestingly, *phnB* and the gene coding for its regulator mvfR, both involved in anthranilate biosynthesis, were upregulated in PAH (Fig. 5C).

### PAH exhibits different biological features

In the infection niche, bacteria are exposed to reactive oxygen species derived from the immune system response and antibiotic treatment (Helaine et al., 2024). Therefore, we tested resistance of PAH to oxidative stress compared to the reference strain PAO1 and the *hmgA\** strain. Oxidative stress resistance was evaluated using a disc diffusion assay with H_2_O_2_ and paraquat. The results revealed that the PAH strain exhibited an intermediate level of resistance compared to PAO1 and the pyomelanin-producing PAO1 *hmgA\** strains (Fig. 6A). When the assay was repeated in minimal medium, the addition of tryptophan notably enhanced the oxidative stress resistance of PAH, with inhibition zones comparable to those observed in LB medium. In contrast, tryptophan supplementation had no appreciable effect on PAO1, indicating a strain specific mechanism of oxidative stress tolerance in PAH. We also tested cytotoxicity in A549 cells using the same concentration of the pigment from PAH and pyomelanin from *hmgA\** strain. While the latter exhibited a half-lethal dose (LD_50_) of 1.33 ± 0.33 mg/mL, while the pigment from PAH showed an LD_50_ of 2.24 ± 0.17 mg/mL. Interestingly, in LB, the proportion of pyo-mel in PAH corresponded to 47% of the total pigment, which was consistent with the differences in the LD_50_, suggesting that the main cytotoxicity was provoked by the soluble fraction of PAH pigment. Along the same line, we evaluated the cytokine response in A549 cells. IL-6 production was nearly undetectable (data not shown), while IL-8 levels increased upon melanin stimulation. The melanin from the PAH strain induced a significantly higher IL-8 response than the pyomelanin derived from the PAO1 *hmgA\** strain (Fig 6B).

**Figure 6.**
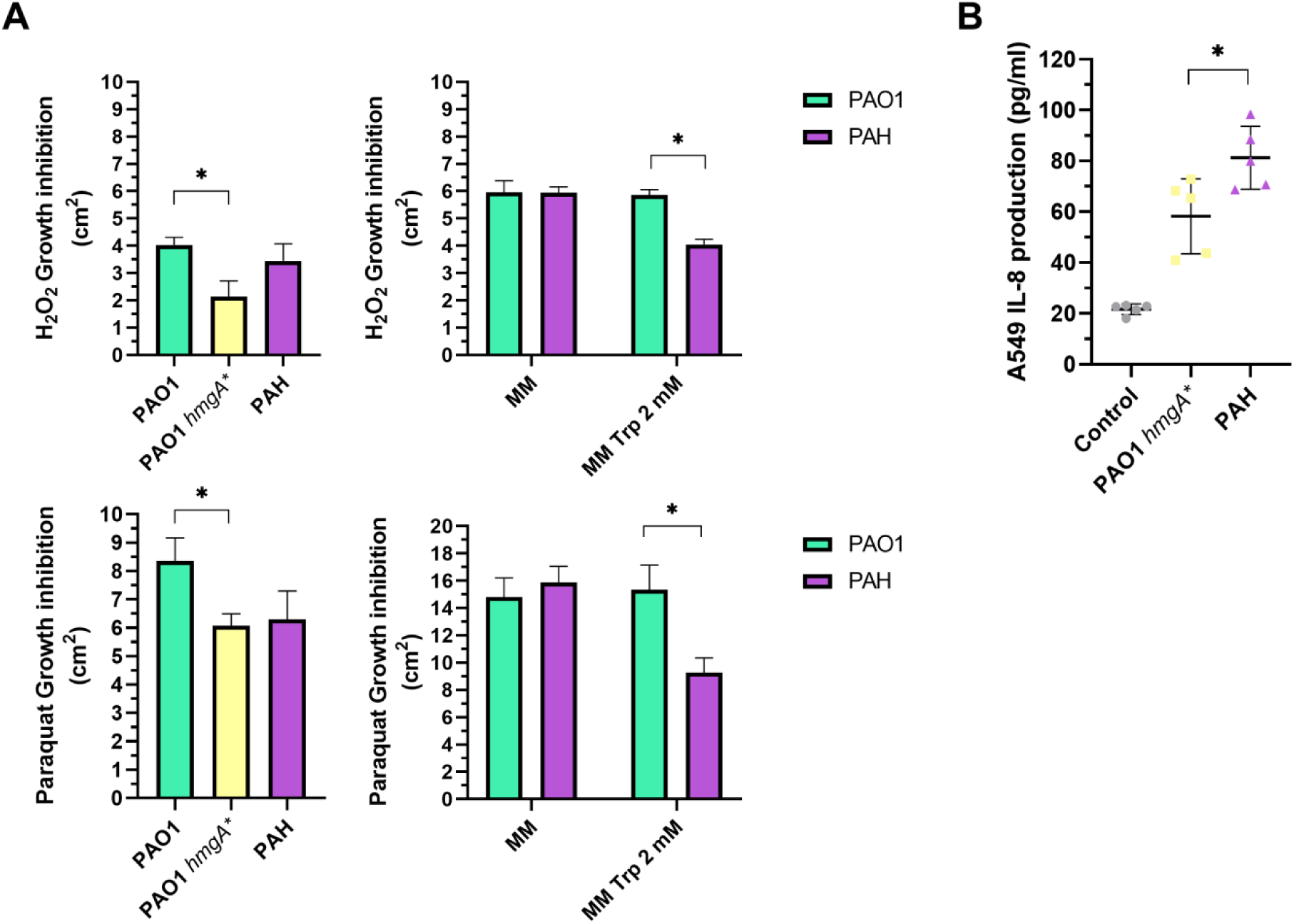
Oxidative stress response and cytokine induction assays. (A) Growth inhibition of *P. aeruginosa* strains in disk diffusion assays with H₂ O₂ and paraquat. Experiments were performed in LB medium for PAO1, PAO1 *hmgA*\*, and PAH, and in minimal medium with or without tryptophan supplementation for PAO1 and PAH. Data represent the mean of four independent replicates. (B) IL-8 cytokine production by A549 cells in response to stimulation with 0.015 mg/mL of purified melanin from PAO1 *hmgA*\* and PAH. No detectable production of IL-6, IL-10, and TNF-α was observed under the tested conditions. Data represent the mean of four independent experiments performed in triplicates. * represents significant differences, P<0.05.

## Discussion

*Pseudomonas* genus comprises a wide spectrum of bacterial species, from Antarctic isolates to pathogenic species causing severe infections (Saati-Santamaría et al., 2022). *P. aeruginosa* is an opportunistic human pathogen, and it is an important cause of morbidity and mortality in hospitalized patients or in those suffering from chronic lung diseases like CF.

PAH was isolated from a child with chronic CF infection (Robaldi et al., 2025, MedRxiv). During the following year, *P. aeruginosa* isolates showing the same phenotypic profile as PAH were repeatedly recovered from this patient, suggesting an adaptation to the CF airway niche. CF airways are shaped by chronic inflammation, areas of bronchiectasis with low oxygen tension, and multiple alterations driven by therapeutic interventions (Van den Bossche, 2021). Altogether, these conditions create a favorable niche for chronic infections (Martin et al., 2023). Despite *P. aeruginosa* carbon and nitrogen metabolic versatility, some nutrients like iron are still essential for biogenesis of cytochromes and Fe-S proteins, which allow bacterial survival and growth. Although iron is abundant on Earth, this metal is poorly available in the presence of oxygen due to readily oxidation to the ferric (Fe^3+^) state, which precipitates in an insoluble form at physiological pH (Murdoch et al., 2022). Inside the host, bacteria must cope with iron deprivation due to nutritional immunity, which withholds essential metals like iron, zinc, and copper (Healy et al., 2021). In *P. aeruginosa*, iron acquisition under limiting conditions relies primarily on the secretion of high-affinity siderophores known as pyoverdines. These compounds chelate Fe³⁺ in the extracellular environment and form soluble complexes that are specifically recognized by outer membrane receptors (Noinaj et al., 2010; Schalk & Guillon, 2013). Once internalized, iron is released through reduction to its ferrous form (Fe²⁺), enabling its utilization in cellular metabolism. Pyoverdine displays a higher affinity for iron than other siderophores such as pyochelin, allowing *P. aeruginosa* to extract iron even from host proteins like transferrin and ferritin (Wolz et al., 1994).

This high-affinity system is crucial for the establishment of acute infections, as evidenced by the upregulation of pyoverdine biosynthetic genes and the attenuated virulence of mutants lacking pyoverdine (Takase et al., 2000; Meyer et al., 1996; Damron et al., 2016). In PAH, the pyoverdin biosynthetic operons and transporters for exogenous pyoverdins presented several frameshifts and other deleterious mutations, resulting in low or no expression. Leinweber et al., (2018) studied pyoverdin production in the presence of different iron concentrations, showing that at intermediate concentrations (0.5 μM), pyoverdin is still important; while ―replete‖ iron concentrations (20 µM) lead to *pdvA* repression by Fur. Moreover, pyoverdins are essential in a medium with apo-transferrin, simulating host iron restrictive conditions (Leinweber et al., 2024, 2018; Meyer et al., 1996). Several studies have shown that pyomelanin mediates ferric to ferrous iron reduction, making it bioavailable under iron-limiting conditions (Turick et al., 2002, 2003). Moreover, in pathogenic species, such as *Legionella pneumophila*, melanin can also effectively reduce Fe^+3^ to Fe^+2^ and release iron from ferritin and transferrin (Zheng et al., 2013).

Furthermore, by scavenging reactive oxygen and nitrogen species (ROS and RNS), melanin enhances resistance to oxidative damage generated by host immune defenses and can modulate these responses (Keith et al., 2007; Ahmad et al., 2016; Nosanchuk and Casadevall, 2006; Plonka and Grabaka, 2006). For example, non-melanogenic *Burkholderia cenocepacia* was more sensitive to oxidative stress, showing reduced survival in macrophages (Keith et al., 2007). Accordingly, pyomelanin-producing clinical isolates of *P. aeruginosa* show increased stress resistance and persistence during chronic lung infections (Rodriguez Rojas et al., 2009; Hocquet et al., 2016). PAH showed increased expression of some genes related to redox balance and higher resistance to oxidative agents compared to non-melanogenic *P. aeruginosa*.

PAH melanin exhibited different characteristics, such as the presence of an NaOH insoluble fraction, when compared to traditional pyomelanins from tyrosine catabolism found in several natural *Pseudomonas* mutant strains. The FTIR spectrum of the total PAH melanin showed similarity with that obtained from the tryptophan-derived melanin from *Rubrivivax benzoatilyticus* JA2 (Ahmad et al., 2020); however, stronger absorptions were recorded at 2924, 2854, 1213, and 1040 cm^-1^ in the PAH pigment, indicative of higher proportions of aromatic and C-O from phenol or aliphatic units. The presence of nitrogen-containing structural units could not be completely confirmed by FTIR spectroscopy, since some absorption bands would overlap with oxygen-related bonds (3276 and 1526 cm^-1^) and the signal at 1414 cm^-1^ detected in the melanin from *R. benzoatilyticus* JA2 could not be found in the PAH spectrum. By applying two different NMR approaches and metabolomic analysis, we showed that the insoluble trp-mel pigment presented structures compatible with indolic compounds derived from tryptophan. Nevertheless, the distribution of the polymeric domains in the overall structure remains unclear, since the different aromatic residues could be crosslinked to give an insoluble hetero-polymer of high molecular weight. Alternatively, some polymeric structures of lower molecular weight might coprecipitate by strong non-covalent intermolecular interactions (Liebscher et al., 2013).

Given that PAH lacks functional pyoverdine synthesis, the redox-heterogeneous pigment may provide a compensatory mechanism for local Fe^3+^-capture, retention and reduction to Fe^2+^, anchoring iron near the bacterial surface despite lower binding affinity than canonical siderophores. Moreover, the more compact and less dispersed appearance of the PAH pigment compared with the pyomelanin produced by the PAO1 *hmgA\**, as observed by TEM, suggested a more hydrophobic nature that could favor iron capture.

Based on our experiments, we propose a model (Fig. 7) for pigment production in PAH triggered by impairment of pyoverdin biosynthesis. This ―iron-limited‖ state drives a metabolic reshaping leading to the production of two types of pigments, pyo-mel and trp-mel, using various metabolic intermediates including tryptophan-derived precursors (Fig. 7A). The relationship between tryptophan and trp-mel precursors is evidenced by the decrease in production when the TDO enzyme is inhibited in PAH (Fig. 7B). In *P. aeruginosa*, Fur directly represses the expression of the sigma factor *pvdS* in iron-replete environments (Leoni et al., 1996). When iron is scarce, this repression is relieved, leading to the activation of the *pvd* cluster responsible for pyoverdine biosynthesis (Cornelis, 2009; Reinhart & Oglesby-Sherrouse, 2016). Under these conditions, PAH would maintain Fur in its apo form (Fig. 7A), thus reshaping iron and carbon metabolism (Oglesby-Sherrouse & Vasil, 2013; Djapgne et al., 2018). Since the pyoverdine biosynthetic cluster is impaired in PAH (Fig. 7A), aromatic intermediates cannot be fully directed toward pyoverdin production. Thus, we hypothesize that intermediates like anthranilate are rerouted into alternative metabolic pathways, promoting trp-mel production in PAH (Fig. 7A).

**Figure 7:**
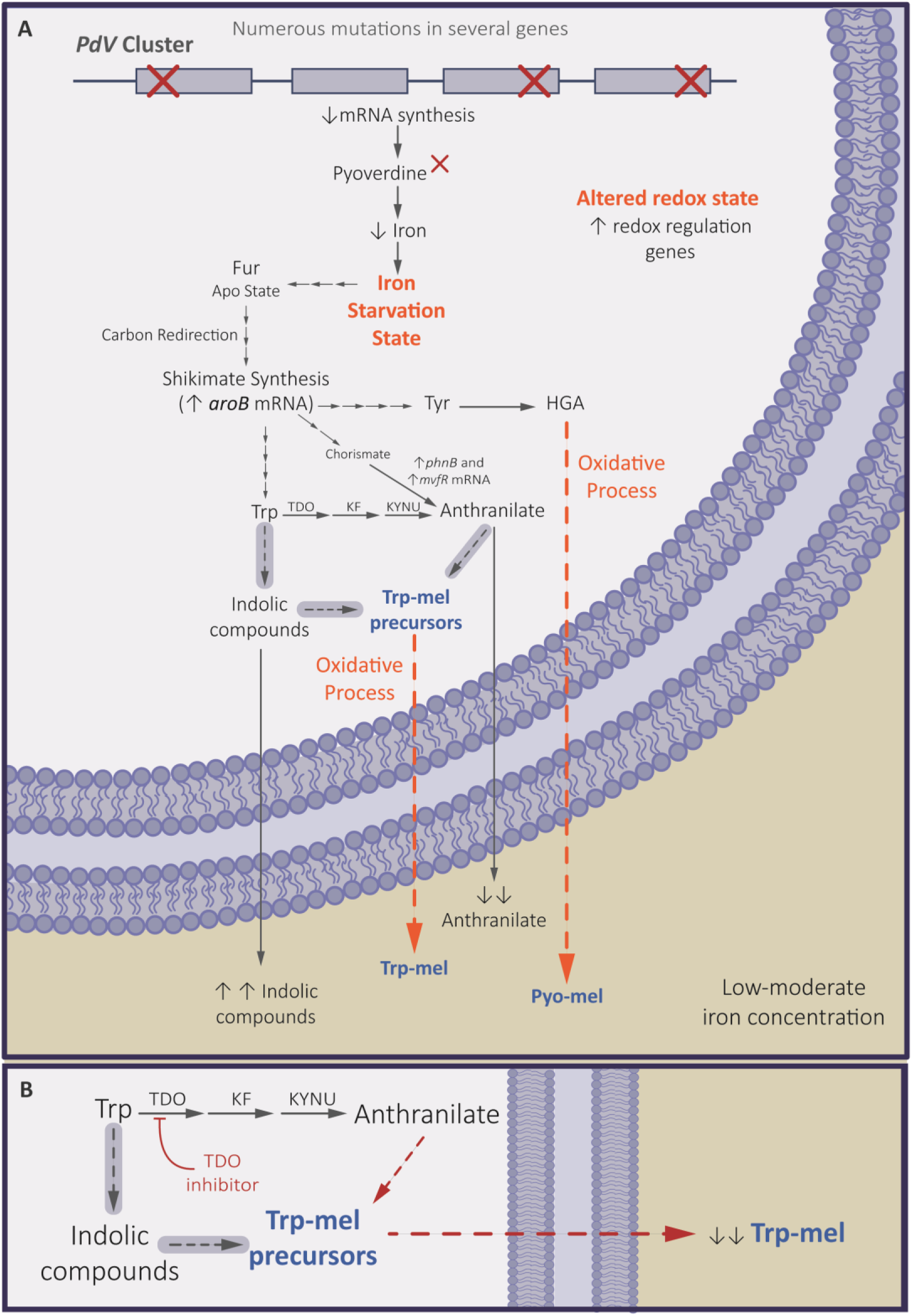
(A) Proposed model for trp-mel and pyo-mel production in *P. aeruginosa* PAH. The iron starvation state due to the impairment of pyoverdin biosynthesis leads to metabolic changes. The shikimate pathway over-feeds the anthranilic and tyrosine biosynthesis branches, increasing the trp-mel and tyr-mel precursors. Crosses indicate genes carrying frameshift mutations or complete gene deletions in PAH. Grey-shaded arrows represent unknown or putative pathways that may contribute to melanin biosynthesis under iron-limited conditions. (B) Effect of TDO inhibition on trp-mel production. Abbreviations: TDO, tryptophan 2,3-dioxygenase; KF, kynurenine formamidase; KYNU, kynureninase.

This process reveals a novel adaptive outcome of iron starvation, coupling disrupted siderophore synthesis with the emergence of melanin production that may contribute to persistence and protection in the cystic fibrosis lung environment.

## Financial statement

This work was supported by funding: To PMT ―Redes Tecnologicas de Alto Impacto, Red REPARA‖ program (Jefatura de Gabinete de Ministros, Argentina), the National Agency for Scientific and Technological Promotion (ANCPyT) PICT-2021-GRFTI-00265, by the (CONICET-PIP-2022); To FG from the Deutsche Forschungsgemeinschaft the Germany‘s Excellence - Strategy—EXC 2124—390838134 ―Controlling Microbes to Fight Infections‖; To MNDA by the Deutscher Akademischer Austauschdienst (DAAD) and the National Secretary of Education of Argentina through the Ale-Arg short-term exchange (Type A; 2024) scholarship; and to AK by Universidad de Buenos Aires (UBACyT 20020220300089BA), CONICET (PIP 1120210100680CO).

## Contributions

MNDA (under PMT, NIL and FG supervision) and NIL, AAK and PMT performed the conceptualization. MNDA, AAK, NIL and PMT performed most of the investigation. MN designed the succinate minimal medium and performed the HPLC experiments. SAR (under PMT supervision) conducted the assembly and in silico analysis of PAH strain. MNDA, NL and EL performed experiments with cell lines and oxidative resistance (under FG supervision). PH conducted MS-RMN experiments and interpreted the data. LP and MA performed the metabolomic experiments and the data analysis. PMC performed the PAH isolation and its identification by MALDI-TOF techniques. MNDA, AAK, NIL and PMT wrote the manuscript. FG, LP, MA, PH, MN reviewed and edited the manuscript. MNDA, NIL, AAK, MA and PMT carried out the visualization. PMT performed mostly the project administration and funding acquisition. AAK, PH, AL, and FG contributed with resources. All authors read and approved the final manuscript.

## Supplementary figures

**Supplementary Figure S1.**
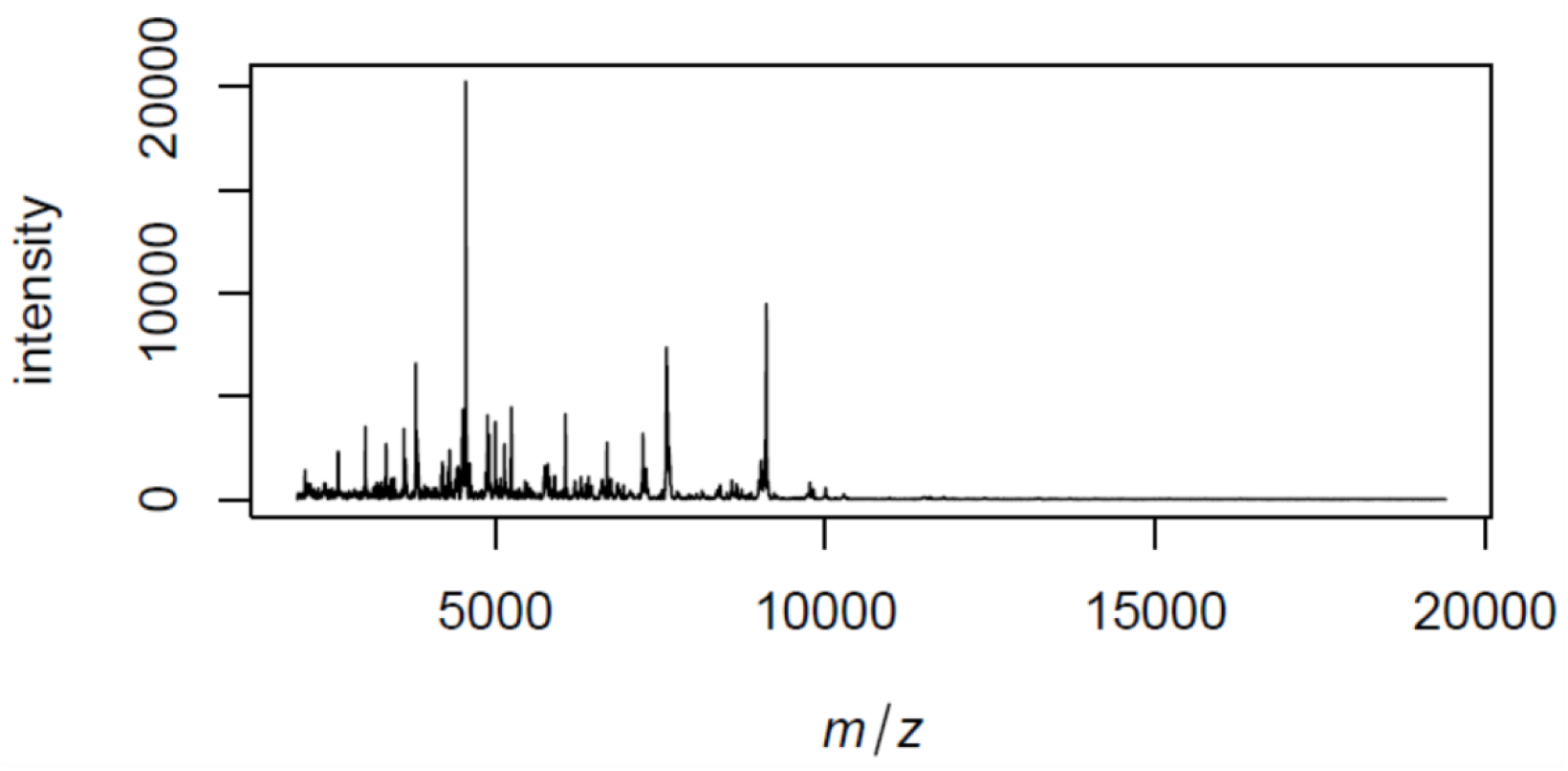
Identification of the PAH isolate as *Pseudomonas aeruginosa* by MALDI-TOF mass spectrometry. Illustrative spectrum obtained as an average of 4 spectra acquired using the VITEK MS platform (*bioMérieux*), based on a Shimadzu Biotech Axima Assurance mass spectrometer (software version 2.9.5.6), under default acquisition and calibration settings provided by the manufacturer. The isolate was identified as *P. aeruginosa* by comparison of the acquired spectrum with VITEK MS in vitro diagnostics (IVD) knowledge database version 3.3. Due to the mucoid nature of the colonies, ethanol-formic acid-acetonitrile extraction using bead beating cell disruption was required to obtain a high-quality spectrum

**Supplementary Figure S2.**
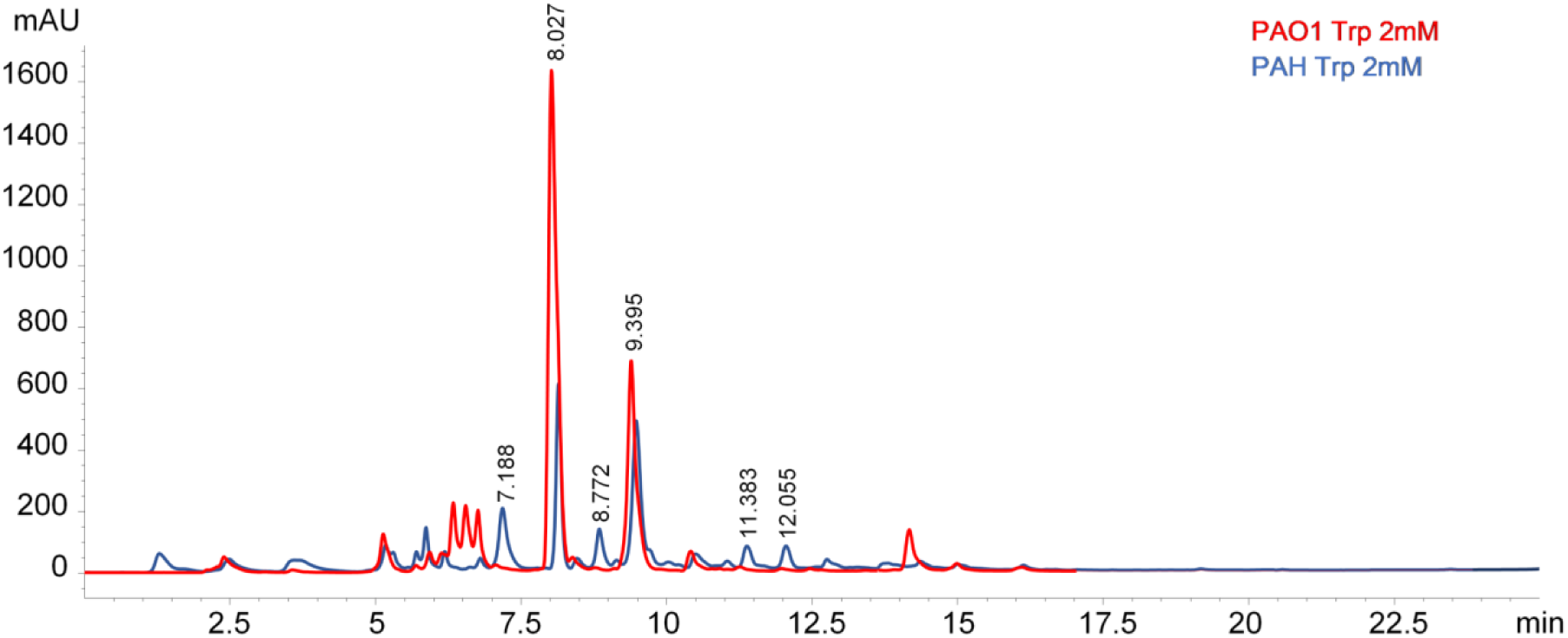
HPLC chromatogram of culture supernatants from PAO1 and PAH grown for 18 h in succinate minimal medium supplemented with tryptophan (2mM). The chromatogram shows the absorbance at 230 nm and is representative of three independent experiments showing similar profiles. The peak detected at 8.02 s retention time was confirmed to correspond to tryptophan by standard comparison.

**Supplementary Figure S3.**
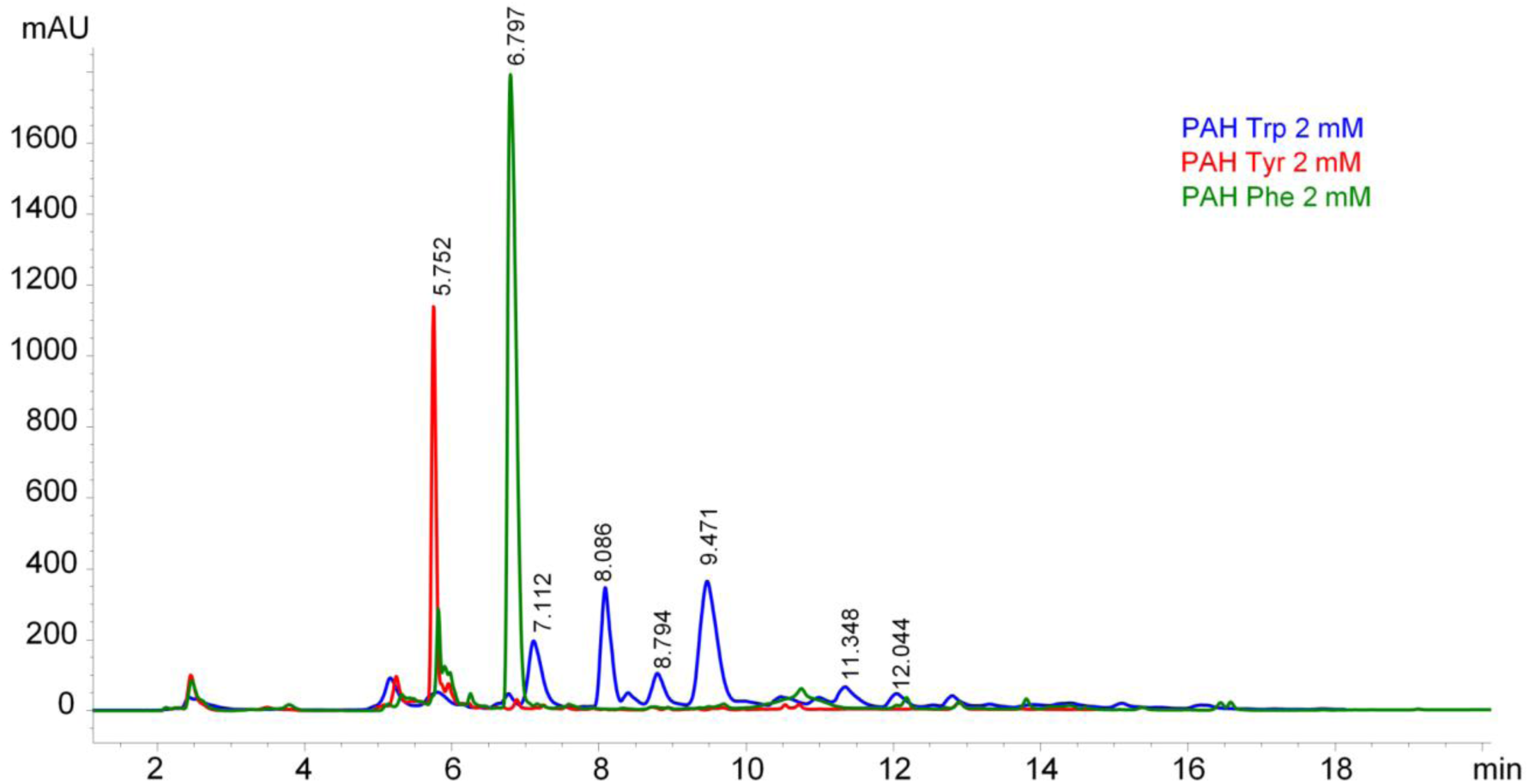
HPLC chromatogram of culture supernatants from PAH grown for 18 h in succinate minimal medium supplemented with tryptophan, tyrosine, or phenylalanine (2 mM). The chromatogram shows the absorbance at 230 nm and is representative of three independent experiments showing similar profiles.

**Supplementary Figure S4.**
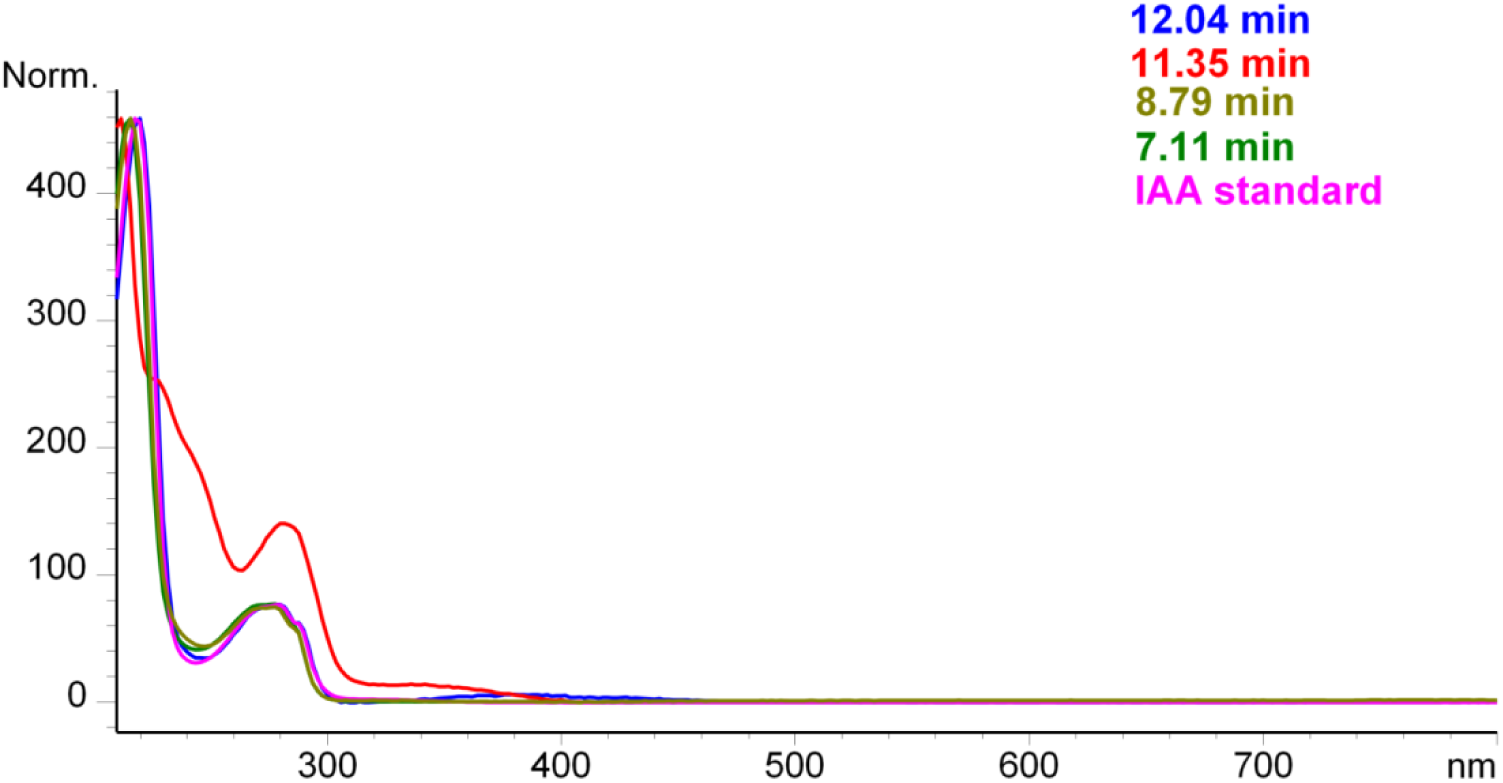
UV-Vis absorption spectra correspond to the peaks shown in Supplementary Figures 2 and 3, with retention times indicated by color. The UV-Vis spectrum of indole-3-acetic acid (IAA) standard (Sigma Aldrich) is included for comparison.

**Supplementary Figure S5.**
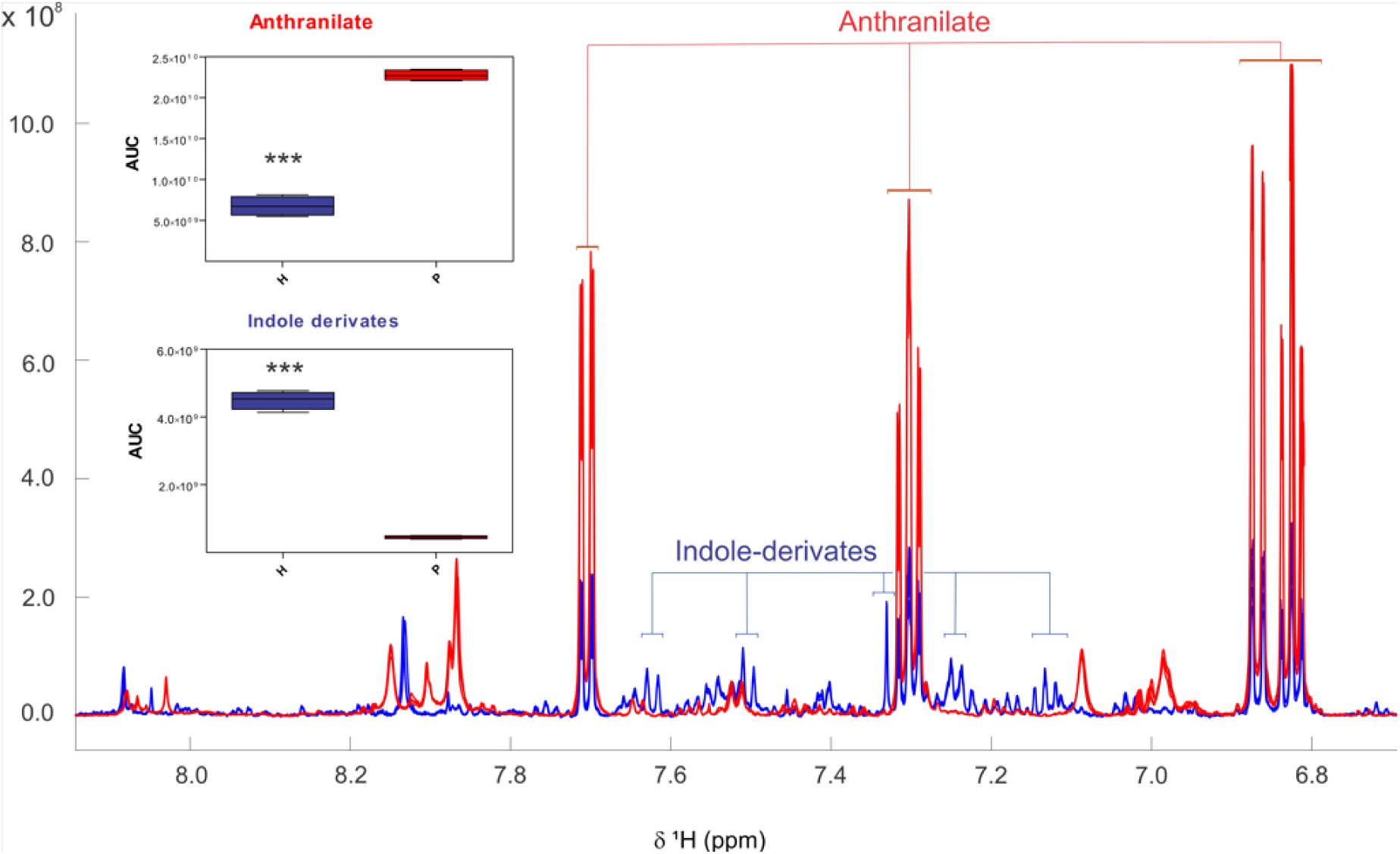
Targeted NMR-based metabolomics of PAO1 and PAH supernatants. Overlaid 1 H-NMR spectra in the aromatic region from four biological replicates of strains PAH (blue) and PAO1 (red) highlight resonances assigned to anthranilate and indole-derived metabolites. The inset depicts the relative area under the curve (AUC) values for the corresponding resonances. Statistical analysis was performed using a non-parametric *t*-test (p<0.0001).

**Supplementary Figure S6.**
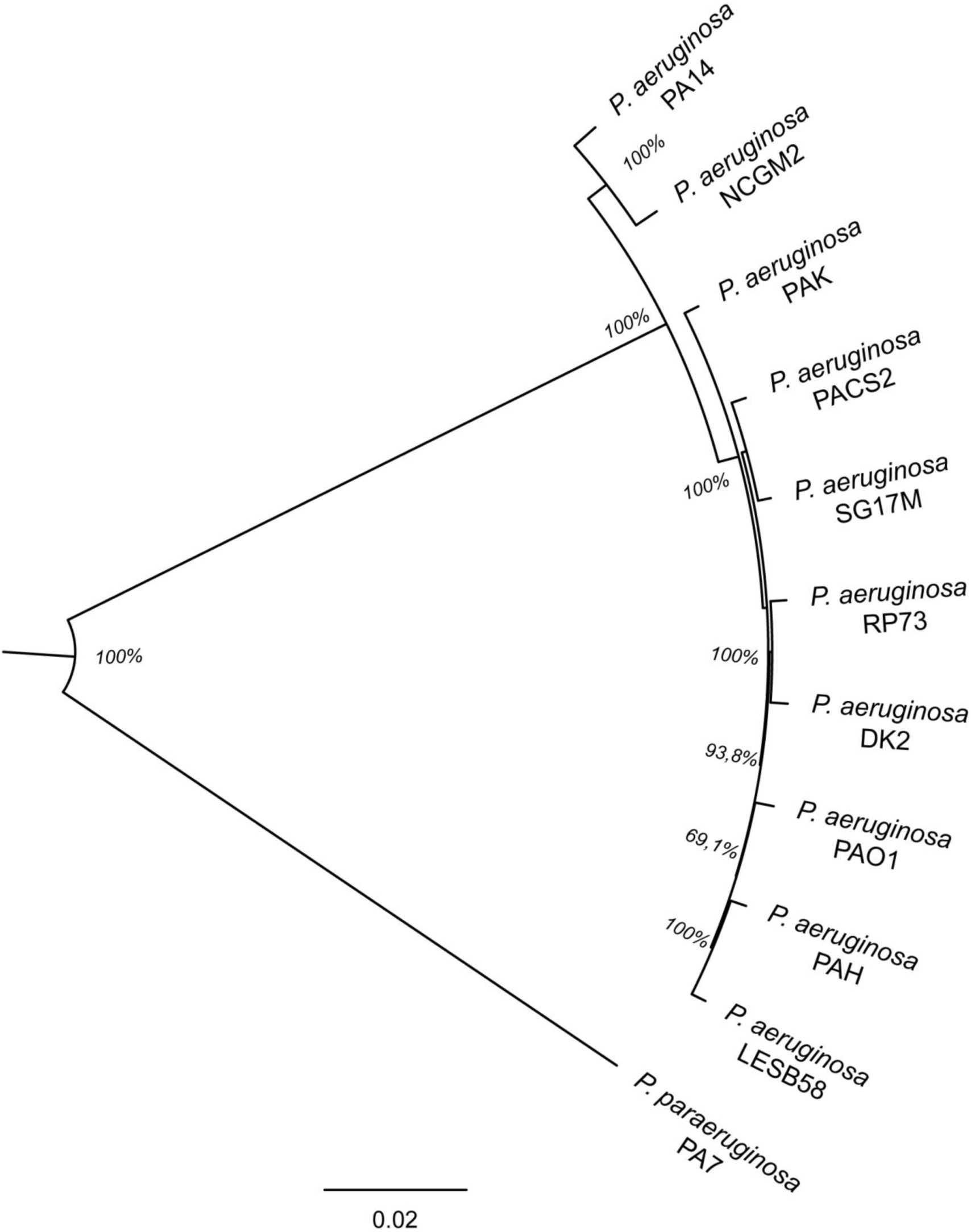
Maximum-likelihood phylogenetic tree based on genome-wide SNP distances for *Pseudomonas aeruginosa* reference strains and the PAH isolate, using PAO1 as the reference genome. *P. paraeruginosa* PA7 was included as an outgroup. Branch lengths indicate substitutions per site (scale bar = 0.02). Numbers on branches show node support values, expressed as the percentage of times that each split was recovered among the 1,000 replicate trees and were assessed using *UFBoot2* (1,000 ultrafast bootstrap replicates) and SH-aLRT (1,000 replicates). The full alignment was exported from *Parsnp* and analyzed with *IQ-TREE2*.

**Supplementary Table S1.** Sequence comparison of genes involved in pigment production and other relevant physiological traits in *P. aeruginosa* PAO1 and PAH strains. Single nucleotide polymorphism (SNP) analysis: variations in single DNA base pairs were classified using a color code: LOW (silent or synonymous mutations, in green), MODERATE (missense mutations, in yellow), HIGH (frameshifts, premature stop codons, insertions, etc., in pink), while positions with no changes compared to the PAO1 homolog were shown in white.

| Gene | PAO1 | aa id. (%) | SNPs |
| --- | --- | --- | --- |
| ***Tryptophan degradation and production of indolic compounds*** |  |  |  |
| *kynA/tdo* | PA2579 | 100 |  |
| *kynR* | PA2082 | 100 |  |
| *kynB* | PA2081 | 100 |  |
| *kynU* | PA2080 | 100 |  |
| *antR* | PA2511 | 99 |  |
| *antA* | PA2512 | 100 |  |
| *antB* | PA2513 | 99 |  |
| *antC* | PA2514 | 99 |  |
| *catA* | PA2507 | 100 |  |
| *catB* | PA2509 | 99 |  |
| *catC* | PA2508 | 100 |  |
| *catR* | PA2510 | 100 |  |
| *amiE'* | PA3797 | 100 |  |
| *amiE* | PA3366 | 99 |  |
| *amiB* | PA3365 | 99 |  |
| *amiC* | PA3364 | 99 |  |
| *amiR* | PA3363 | 99 |  |
| *amiS* | PA3362 | 100 |  |
| ***Tyrosine-phenylalanine catabolism*** |  |  |  |
| *phhR/tyrR* | PA0873 | 99 |  |
| *phhA* | PA0872 | 99 |  |
| *phhB* | PA0871 | 100 |  |
| *phhC/tyrB* | PA0870 | 99 |  |
| *hpd/hppD* | PA0865 | 100 |  |
| *hmgR* | PA2010 | 99 |  |
| *hmgA* | PA2009 | 100 |  |
| *maiA* | PA2007 | 99 |  |
| *fahA* | PA2008 | 99 |  |
| ***Transporters of HGA, membrane and hypothetical proteins*** |  |  |  |
| *hatA/yrbF* | PA4456 | 100 |  |
| *hatB/yrbE* | PA4455 | 100 |  |
| *blc* | PA5107 | 99 |  |
| *kup* | PA0917 | 99 |  |
| - | PA2963 | 100 |  |
| - | PA1190 | 100 |  |
| **Transcription and regulation** |  |  |  |
| *cbrB* | PA4726 | 99 |  |
| *ntrC* | PA5125 | 99 |  |
| *crcB* | PA4383 | 100 |  |
| *rpoN* | PA4462 | 100 |  |
| *mvaU* | PA2667 | 100 |  |
| - | PA4112 | 99 |  |
| **Pyoverdine synthesis** |  |  |  |
| *pvdS* | PA2426 | 100 |  |
| *pvdG* | PA2425 | 99 |  |
| *pvdL* | PA2424 | 99 |  |
| *pvdH* | PA2413 | 100 |  |
| - | PA2412 | 100 |  |
| - | PA2411 | 99 |  |
| *fpvF/ABC3* | PA2410 | 99 |  |
| *fpvE/ABC2* | PA2409 | 100 |  |
| *fpvD/ABC1* | PA2408 | 99 |  |
| *fpvC* | PA2407 | 99 |  |
| *fpvK* | PA2406 | 99 |  |
| *fpvJ* | PA2405 | 100 |  |
| *fpvH* | PA2404 | 96 |  |
| *fpvG* | PA2403 | 84 |  |
| *pvdI* | PA2402 |  |  |
| *pvdJ* | PA2400 | 47 |  |
| *pvdD* | PA2399 |  |  |
| *fpvA* | PA2398 |  |  |
| *pvdE* | PA2397 | 57 |  |
| *pvdF* | PA2396 | 95 |  |
| *pvdO* | PA2395 | 97 |  |
| *pvdN* | PA2394 | 93 |  |
| *pvdM* | PA2393 | 87 |  |
| *pvdP* | PA2392 | 88 |  |
| *ompQ* | PA2391 | 99 |  |
| *pvdT* | PA2390 | 99 |  |
| *pvdR* | PA2389 | 99 |  |
| *fpvR* | PA2388 | 99 |  |
| *fpvI* | PA2387 | 100 |  |
| *pvdA* | PA2386 | 99 |  |
| *pvdQ* | PA2385 | 99 |  |
| **Ferrous iron (Fe<sup>2+</sup>) uptake genes** |  |  |  |
| *feoB* | PA4358 | 99 |  |
| *feoA* | PA4359 | 100 |  |
| *feoC* | Absent | 98 |  |
| ***Fe<sup>3+</sup>–siderophore (pyochelin) uptake*** |  |  |  |
| *fptA* | PA4221 | 99 |  |
| *pchG* | PA4224 | 99 |  |
| *pchF/H* | PA4225 | 99 |  |
| *pchE/I* | PA4226 | 99 |  |
| *pchR* | PA4227 | 100 |  |
| *pchD* | PA4228 | 99 |  |
| *pchC* | PA4229 | 99 |  |
| *pchB* | PA4230 | 99 |  |
| *pchA* | PA4231 | 99 |  |
| ***Uptake of xenosiderophores (siderophores from other bacteria)*** |  |  |  |
| *fiuA* | PA0470 | 99 |  |
| *pfeA* | PA2688 | 99 |  |
| *foxA* | PA2466 | 100 |  |
| ***Iron uptake from heme/hemoglobin*** |  |  |  |
| *phuR* | PA4710 | 100 |  |
| *hasE* | PA3405 | 99 |  |
| *hasD* | PA3406 | 97 |  |
| *hasA* | PA3407 | 99 |  |
| *hasR* | PA3408 | 99 |  |
| *hasS* | PA3409 | 98 |  |
| ***Regulation of iron homeostasis*** |  |  |  |
| *fur* | PA4764 | 100 |  |
| *fiuR* | PA0471 | 100 |  |
| *pfeR* | PA2696 | 98 |  |
| ***TonB-dependent transporters related to iron*** |  |  |  |
| *pirA* | PA0931 | 99 |  |
| *piuA* | PA4514 | 99 |  |
| ***Aminoacid metabolism*** |  |  |  |
| *aroB* | PA5038 | 99 |  |
| *cysE* | PA3816 | 99 |  |
| *pyrB* | PA0402 | 100 |  |
| *dapA* | PA1010 | 100 |  |
| *dapB* | PA4759 | 100 |  |
| *argE* | PA5206 | 99 |  |
| *thrC* | PA3735 | 99 |  |
| *iscS* | PA3814 | 100 |  |

